# From Development to Regeneration: Insights into Flight Muscle Adaptations from Bat Muscle Cell Lines

**DOI:** 10.1101/2025.07.01.662643

**Authors:** Fengyan Deng, Valentina Peña, Pedro Morales-Sosa, Andrea Bernal-Rivera, Bowen Yang, Shengping Huang, Sonia Ghosh, Maria Katt, Luciana Castellano, Cindy Maddera, Zulin Yu, Nicolas Rohner, Chongbei Zhao, Jasmin Camacho

## Abstract

Skeletal muscle regeneration depends on muscle stem cells, which give rise to myoblasts that drive muscle growth, repair, and maintenance. In bats—the only mammals capable of powered flight—these processes must also sustain contractile performance under extreme mechanical and metabolic stress. However, the cellular and molecular mechanisms underlying bat muscle physiology remain largely unknown. To enable mechanistic investigation of these traits (Graphical Abstract), we established the first myoblast cell lines from the pectoralis muscle of *Pteronotus mesoamericanus*, a highly maneuverable aerial insectivore. Using both spontaneous immortalization and exogenous hTERT/CDK4 overexpression, we generated two stable cell lines that retain proliferative capacity and differentiate into contractile myotubes. These cells exhibit frequent spontaneous contractions, suggesting robust functional integrity at the neuromuscular junction. In parallel, we performed transcriptomic and metabolic profiling of native pectoralis tissue to define molecular programs supporting muscle specialization. Gene expression analyses revealed enriched pathways for muscle metabolism, development, and regeneration, highlighting the supporting roles in tissue maintenance and repair. Consistent with this profile, the flight muscle is triglyceride-rich, which serves as an important fuel source for energetically demanding processes, including muscle contraction and cellular recovery. Integration of transcriptomic and metabolic data identified three key metabolic modules—glucose utilization, lipid handling, and nutrient signaling—that likely coordinate ATP production and support metabolic flexibility. Together, these complementary tools and datasets provide the first in vitro platform for investigating bat muscle research, enabling direct exploration of muscle regeneration, metabolic resilience, and evolutionary physiology.

**Graphical Abstract:** Establishment of bat muscle cell cultures from the Mesoamerican mustached bat (*P. mesoamericnus*) provides an in vitro platform to investigate muscle regeneration and flight muscle biology. The pectoralis major muscle was isolated to generate primary myoblast cultures, which were expanded and immortalized using hTERT/CDK4. The resulting myoblast lines retain proliferative and differentiation capacity. RNA sequencing of native pectoralis muscle tissue revealed molecular signatures of myogenic regulation, stress resilience, and tissue remodeling, supporting the relevance of these in vitro models for studying muscle maintenance and regenerative mechanisms.

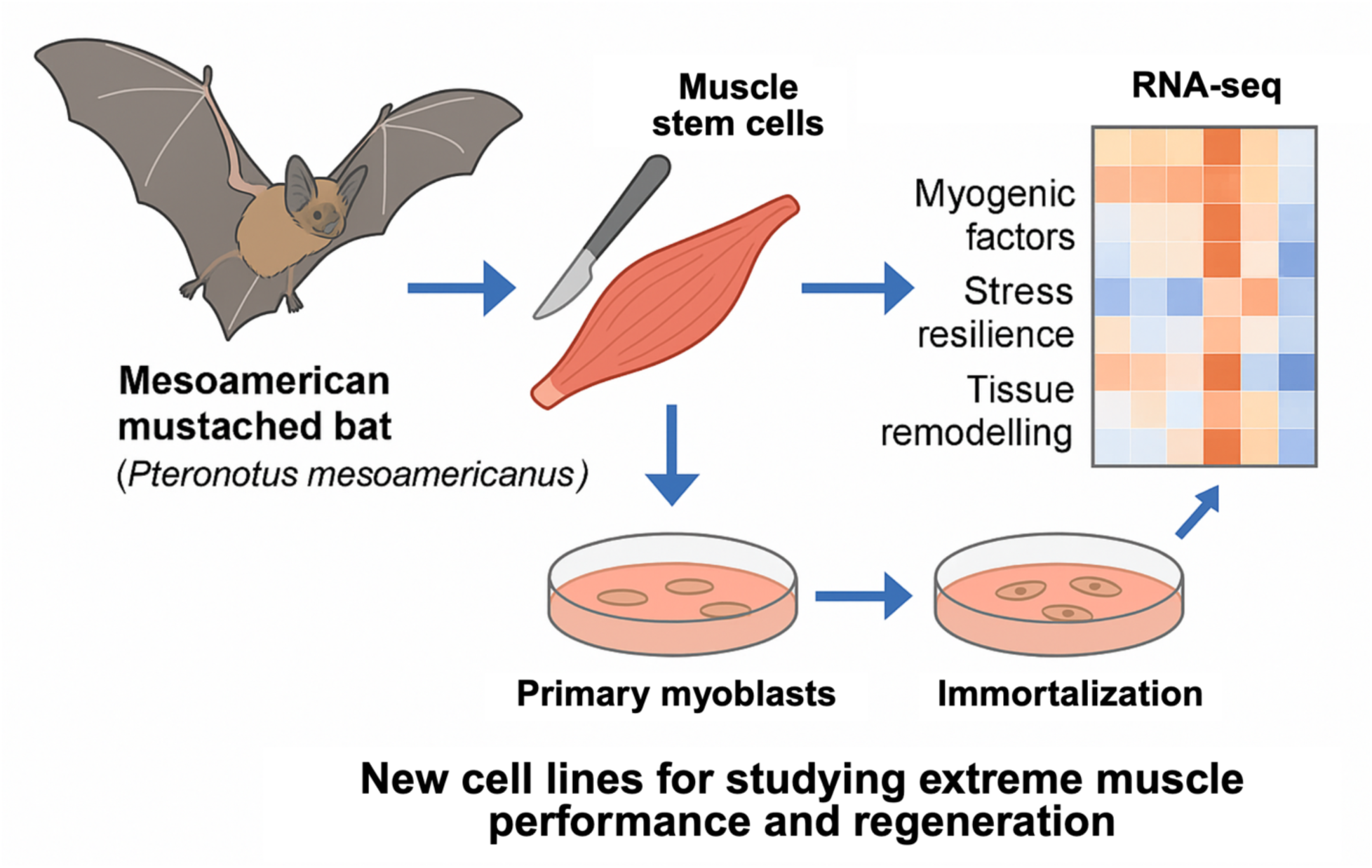

## **1.** Introduction

Understanding how skeletal muscle regenerates and maintains function under stress is critical for addressing muscle loss, injury, and degenerative diseases. Myoblasts, the progenitor cells of skeletal muscle, play central roles in both muscle development and repair [1–3]. While most models in muscle biology rely on human and rodent cells, expanding to non-traditional species offers an opportunity to uncover lineage-specific adaptations of regeneration and functional resilience, especially in mammals with extreme physiological traits (Table 1).

**Table 1.** Summary of Established Myoblast Cell Lines Across Vertebrate Species. A comparative overview of established myoblast and muscle-derived cell lines from various vertebrate species, including their typical research uses. These cell lines have been applied in diverse fields such as muscle physiology, regenerative biology, disease modeling, metabolic research, aquaculture, and gene therapy development. This table highlights the broad utility of muscle cell models and emphasizes the gap in non-traditional systems such as bats, which possess unique physiological traits including flight, hibernation, longevity, and enhanced regenerative capacity.

| Species/Source | Cell Line Name | Research Applications | References |
| --- | --- | --- | --- |
| Rat | L6 | Muscle metabolism, insulin signaling, hypertrophy | [72, 87] |
| Mouse | C2C12 | Myogenesis, muscle regeneration, gene expression |  |
| Human (Myotonic Dystrophy) | Myotonic Dystrophy Cell Lines | Myotonic dystrophy disease modeling, therapy screening | [46, 47] |
| Dog (Myok9) | Myok9 | Canine muscle disease and gene therapy studies | [88] |
| Dog (Dystrophic) | Dystrophic Myoblast Cell Lines | Muscular dystrophy modeling, therapy testing | [49] |
| Human (Healthy Muscle) | Human Myoblast Cell Line | Muscle aging, physiology, regenerative medicine | [36] |
| Grass Carp | CIM | Aquatic muscle growth, stress physiology | [74] |
| Crab-eating macaque | NHP iPAX7 | Myotube differentiation and muscle regeneration | [89] |
| Japanese Eel | JEM1129 | Fish muscle development, aquaculture research | [90] |
| Brown-Marbled Grouper | EfMS | Fish muscle stem cell and regenerative studies | [51] |
| Chicken | chTERT-Myoblasts / Primary Chicken Myoblasts | Avian muscle biology, myogenesis, gene studies | [91] |
| Cuvier's beaked whale | pSV3neo myoblast cell line | Extreme hypoxia, fasting, and deep-diving | [92] |
| Bat | iBatM-Pmeso-S1<br>iBatM-Pmeso-TC1 | Flight, muscle endurance, muscle stem cell and regeneration, metabolic resilience | This paper |

The evolution of powered flight in bats represents one of the most dramatic physiological transformations in mammalian history. As the only mammals capable of flight, bats underwent rapid morphological and functional innovations [4, 5]. A hallmark of this transition is the specialization of skeletal muscle—not only for force production, but for sustaining the intense and continuous energy output during flight . Although the developmental changes underlying bat wing morphology—including elongation of the digits, formation of the wing membrane, and specialization of flight muscles—have been partially characterized [7–11], yet the molecular and metabolic programs that support the evolution of flight, particularly those related to muscle regeneration and energy balance, remain poorly understood [12–14].

Bats represent a diverse group of mammals known for their distinctive physiological traits, including exceptional longevity without age-related reproductive decline [15, 16], low cancer incidence [17, 18], dampened inflammatory responses [19–22], viral resistance [22], rapid wound healing, enhanced DNA repair [24], and the capacity to enter hibernation or torpor . These traits suggest that bats possess robust mechanisms for tissue protection and long-term physiological resilience. This makes them a compelling model for studying muscle biology, particularly in the context of regeneration, mitochondrial oxidative stress, and aging [15, 16]. Their muscles must remain functional under intense workloads, recover efficiently from damage, and resist age-related decline . Recognizing this potential, establishing in vitro bat muscle models represents a key step toward uncovering the molecular mechanisms underlying their exceptional stress-tolerant physiology [27–32].

Culturing primary myoblasts poses a major challenge due to their limited lifespan in vitro . Mammalian myoblasts rapidly lose their stemness shortly after isolation and rarely maintain it beyond 10 passages as a result of cellular senescence [34, 35]. Senescence is primarily driven by two key mechanisms [36, 37]. First, progressive telomere shortening occurs with each cell division until they reach a critical length that activates the p53 pathway and leads to cell senescence [38–41]. Telomerase reverse transcriptase (TERT), catalytic subunit of the telomerase complex, maintains telomere length and delays senescence. Its expression into primary cells has been shown to extend lifespan and preserve proliferative potential [42, 43]. Second, senescence is mediated by the cyclin-dependent kinase inhibitor (CDKI), p16, which inhibits CDK4/6 kinases activity and causes G1 cell cycle arrest . Overexpression of CDK4 can bypass this arrest, maintaining cell proliferation while preserving normal cell functions [45–47]. Together, co-expression of TERT and CDK4 has enabled the successful immortalization of myoblasts in several species [36, 45, 46, 48–51]. Despite this progress in immortalizing myoblasts from multiple species, no existing models capture the extreme physiological performance and tissue resilience observed in bats (Table 1).

Although advances in comparative genomics in bats, functional models for mechanistic investigation of bat muscle biology remain absent. This gap limits our ability to study the cellular and molecular basis of muscle performance, regeneration, and stress resilience. To address this, we selected *Pteronotus mesoamericanus*, a Neotropical aerial insectivorous bat and member of the diverse Noctilionoidea superfamily, which represents 16% of global bat diversity. This species is capable of high-speed, high-endurance flight [53, 54], and belongs to a subset of *Pteronotus* species—along with *P. parnellii* and *P. personatus*—that have evolved Doppler-sensitive sonar, a specialized form of echolocation requiring rapid neuromuscular precision [55, 56]. Genomic studies have further identified signatures of molecular adaptation in this species across pathways related to stress tolerance, longevity, and immune modulation [57, 58], positioning it as an ideal system for exploring the physiological demands of powered flight. By integrating traits essential for flight and acoustic precision, both of which require finely tuned neuromuscular control and sustained metabolic output, make *P. mesoamericanus* a compelling yet underutilized model for advancing studies of skeletal muscle specialization under extreme physiological stress (Fig. 1).

**Figure 1.**
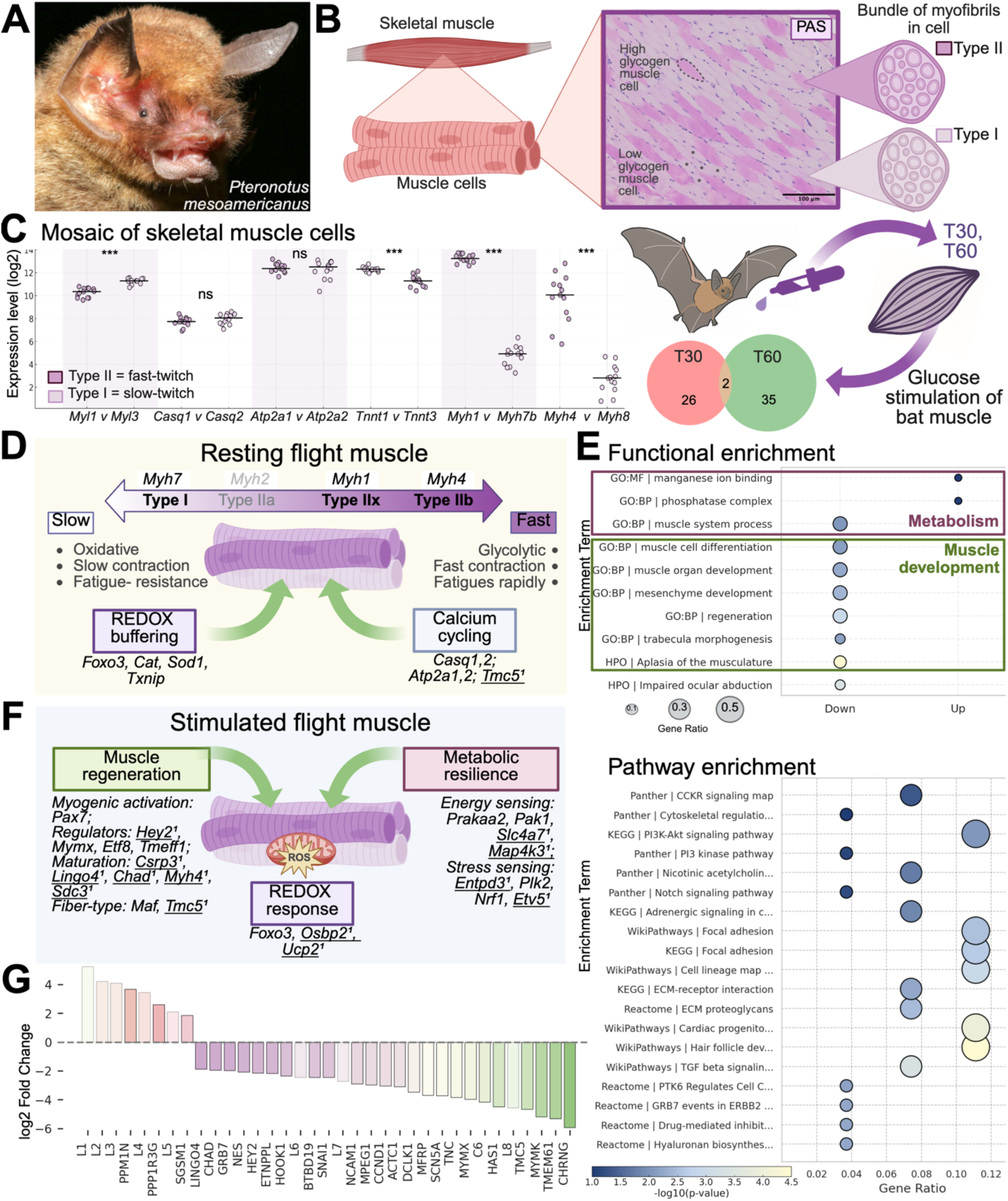
Pteronotus bat as a model for flight muscle specialization and stress resilience. A. Photo of *P. mesoamericanus*, a Neotropical mustached bat specialized for high-speed, high-endurance flight. B. Schematic of skeletal muscle highlighting a mosaic of fast-twitch (Type II) and slow-twitch (Type I) muscle fibers. Periodic Acid-Schiff (PAS) staining reveals high and low glycogen-containing fibers across muscle cells. Skeletal muscle tissue (pectoralis) was sampled at three timepoints: control animals (n=4) represent the baseline state (T0), while glucose-fed animals were sampled 30 (T30; n=3) and 60 (T60; n=5) minutes after oral glucose intake. RNA-seq was performed on flight muscle to identify differentially expressed genes (DEGs) in response to glucose. The Venn diagram shows DEGs between timepoints at 30 minutes (T30 vs T0) and 60 minutes (T60 vs T0). C. Expression of muscle fiber-type genes across 13 individuals, comparing fast-twitch (dark purple) and slow-twitch (light purple) paralogs. Candidate genes include myosin heavy chain (*Myh1*, *Myh4*, *Myh7*, *Myh8*), myosin light chain (*Myl1*, *Myl3*), calsequestrin (*Casq1, Casq2*), calcium ATPases (*Atp2a1, Atp2a2*) and troponin T (*Tnnt3*, *Tnnt1*). Horizontal line represents the median; statistical comparisons were performed using paired t-tests, *** p < 0.001. D. Summary diagram of resting flight muscle gene expression. The pectoralis has a mosaic of fiber types ranging from slow-twitch oxidative (Type I, *Myh7*) to fast-twitch glycolytic fibers (Type IIb, *Myh4*), with intermediate fast-twitch oxidative-glycolytic fibers (Type IIx, *Myh1*). At rest, flight muscle is enriched for gene programs involved in redox buffering (*Foxo3*, *Cat, Sod1, Txnip*) and calcium handling (*Casq1/2*, *Atp2a1/2, <u>Tmc5</u>*^†^). <u>^†^ indicates DEGs with p<0.003 and log_2_ fold change >1.5.</u> E. Enrichment dot plots based on gene set enrichment analysis for T30 vs T60. The top functional terms were associated with muscle metabolism and development. The circle size reflects the gene ratio; color indicates -log10(p-value). Directionality is shown relative to T30: Up = higher in T60, Down = lower in T60. For mechanistic insights, the top 5 enriched terms per pathway-level source (KEGG, Reactome, Panther, WikiPathways) are shown. F. Summary diagram of stimulated flight muscle gene expression grouped by functional modules. Regeneration signaling includes subcategories myogenic activation (*Pax7), regulators (Mymx, <u>Hey2^†^</u>, Etf8, Tmeff1),* maturation (*<u>Lingo4^†^</u>, <u>Chad^†^</u>, <u>Csrp3^†^</u>, <u>Myh4^†^</u>*), and fiber-type specification (*Maf, <u>Tmc5^†^</u>).* Metabolic resilience subcategories include energy sensing (*Prkaa2*, *Pak1, <u>Slc4a7^†^</u>*, *<u>Map4k3^†^</u>*) and stress signaling (*<u>Etv5^†^</u>*, *Plk2, Nrf1*). Redox response: *Foxo3*, *<u>Osbp2^†^</u>*, and *<u>Ucp2^†^</u>*. G. Log_2_ fold-change in expression of genes following glucose stimulation, T60 relative to T30. All data marked † are significant from RNA-seq differential expression analysis.

In this study, we pursued two complementary goals to advance the study of muscle specialization and resilience in bats. First, we performed RNA sequencing of native pectoralis muscle to investigate gene expression associated with muscle performance and stress response in a high-performance, long-lived bat. This included RNA sequencing of muscle tissue collected at rest and after acute glucose stimulation (5.4g/kg body weight), a physiological proxy for activity-induced metabolic stress. We integrated this with systemic glucose dynamics via glucose tolerance testing (GTT) and biochemical quantification of tissue energy stores (triglycerides and glycogen) to characterize muscle energetics in vivo. Second, to functionally explore these pathways and create an experimental platform for bat muscle biology, we established and characterized two immortalized myoblast cell lines derived from the same species. One line arose through spontaneous immortalization, while the other was engineered via hTERT/CDK4 overexpression to overcome replicative senescence. By combining energy-responsive gene expression profiling in tissue with the development of bat muscle cell lines, our study provides a systems-level view into the evolutionary pressures of flight and a cellular model to dissect molecular mechanisms of metabolic adaptation and regeneration. Together, these complementary approaches—transcriptomic analysis of stimulated flight muscle and the establishment of bat-derived myoblast lines—provide a powerful framework for uncovering the cellular and metabolic adaptations of stress tolerance, metabolic flexibility, and sustained muscle function under extreme physiological demands.

## 2. Materials and Methods

### 2.1 Sample collection

A single adult male, *Pteronotus mesoamericanus* (BZ#3524), was collected for cell line development by J.C. in April 2024 at the Lamanai Archaeological Reserve, Orange Walk District, Belize. Animal capture, handling, and euthanasia followed the American Society of Mammalogists’ guidelines for the ethical use of wild mammals in research (Sikes 2016). All procedures were approved by the Institutional Animal Care and Use Committee at the American Museum of Natural History (AMNHIACUC-20240130) and the Stowers Institute for Medical Research (IACUCLASF2024-170). The bat was captured using ground-level mist nets and transported in an individual cloth bag to the Lamanai Field Research Center. Euthanasia was performed using isoflurane inhalation (<1 mL applied to a cotton ball) followed by cardiac puncture for exsanguination. This method induces rapid unconsciousness and humane death within 1–2 minutes due to the high respiratory rate of bats.

The pectoralis major muscle of *P. mesoamericanus* bat was dissected immediately post-euthanasia and rinsed three times in chilled 1x PBS, followed by obtaining four 5 mm circular biopsies using a sterile biopsy punch (Integra™ 3335) in a sterile petri dish filled with chilled 1x PBS. Two biopsies were flash frozen in liquid nitrogen for downstream RNA-sequencing. The remaining two were briefly rinsed in chilled biopsy media (2% FBS + 1mM EDTA in 1x PBS) then quickly transferred into chilled myoblast isolation media (DMEM/F12 with 15 mM HEPES buffer plus 1x MyoCult^TM^ Expansion Supplement). Each biopsy was placed into a sterile microcentrifuge tube containing 2.0 mL of myoblast media with 0.1% Collagenase type I (Sigma, Cat#: SCR103) and minced with surgical scissors (Fine Science Tools, Cat#: 91408-14) until no visible tissue clumps remained. Tubes were incubated for 30 min in a 37°C water bath. After enzymatic digestion, tissue was mechanically dissociated by gentle trituration using a P1000 pipette. Cell suspensions were passed through a 100 μm cell strainer into a 12-well culture plate. The filtered cells were transferred into sterile microcentrifuge tubes, centrifuged at 350 xg for 5 min at ambient temperature (∼27°C), and the supernatant was discarded. Pelleted cells were resuspended in 2.0 mL of myoblast media with 10% DMSO and transferred to cryovials. Cryovials were placed into a Corning® CoolCell® and cooled (−1°C/min) on dry ice for 1.5 hours before storage and shipment in vapor-phase liquid nitrogen. Samples were collected and exported under Belize Forest Department permit FD/WL/7/24 (56) and FD/WL/7/24 (57). Cells were maintained in -80°C until primary cell derivation.

### 2.2 RNA-sequencing

#### 2.2.1 RNA extraction

Skeletal muscle tissue was obtained from male *Pteronotus parnellii* bats (n=13), a species formerly considered conspecific with *P. mesoamericanus* (Clare et al. 2013). Although a distinct species, *P. parnellii* shares key ecological, physiological, and behavioral traits with *P. mesoamericanus*, including large pectoralis muscles for flight. These individuals had previously undergone glucose tolerance testing during field studies in April 2022 [59] collected and exported under Trinidad Forestry Division permit #734. As the primary site of post-prandial glucose uptake and major driver of energy demand during flight, the pectoralis is a key tissue for uncovering metabolic adaptations. Approximately 10 mg of flash-frozen pectoral muscle was homogenized in 0.4 mL of lysis buffer/homogenization solution using a BeadBug 6 microtube homogenizer (Benchmark Scientific) containing zirconium oxide beads, bead size 1.5 mm. Total RNA was extracted following the manufacturer’s protocol for the Maxwell® RSC simplyRNA Tissue Kit (Promega, Cat# AS1340). RNA concentration and integrity were assessed prior to sequencing. Additional details regarding library preparation and sequencing are available in the Supplemental Methods (available at https://www.stowers.org/research/publications/LIBPB-2563).

#### 2.2.2 RNA Sequencing and Read Processing

Total RNA was submitted for poly-A–selected, stranded RNA sequencing using the NEBNext® Ultra™ II RNA Library Prep Kit. Libraries were sequenced on the AVITI-75 platform (Element Biosciences) using paired-end reads. Ribo-depleted, poly-A–enriched libraries were sequenced with temporal resolution to assess differential gene expression over time. Raw sequencing reads were demultiplexed using Illumina bcl-convert v3.10.5, allowing up to one mismatch per index. Reads were aligned to the *P. mesoamericanus* genome (UCSC GCF_021234165.1) using the STAR aligner (v2.7.10b) with NCBI_2023 gene model annotations. Transcript abundance was quantified as TPM using RSEM v1.3.1. Initial quality control included Spearman correlation analysis to assess consistency across biological replicates. Sample correlations were used to identify outliers and validate time-point groupings.

#### 2.2.3 Differential Gene Expression Analysis

Orthologous genes were identified using annotations from the NCBI RefSeq database to enable comparison of conserved transcriptional responses to glucose stimulation and facilitating functional interpretation based on better-characterized gene annotations in other species. Downstream analysis was performed in R v4.3.1 using the edgeR package (v3.36.0). Genes with low expression were filtered out, reducing the dataset from 22,050 annotated genes to 12,406 expressed genes. Normalization of read counts and differential expression testing were carried out in edgeR, with DEGs defined as those with an adjusted *p*-value < 0.05 and absolute log_2_ fold change > 1. GO annotations were obtained from NCBI RefSeq and applied to differentially expressed genes (DEGs) within each treatment group. GO enrichment analysis was performed using ClusterProfiler v4.10.0, with enrichment based on DEGs passing thresholds of *p* < 0.05 and absolute fold change > 1.5. Enriched biological processes and cellular components were used to contextualize gene expression patterns related to metabolic adaptation.

#### 2.2.4 Functional Profiling of Metabolically Stimulated Differentially Expressed Genes

To identify pathways relevant to muscle physiology and flight metabolism, we analyzed differentially expressed genes (DEGs) from *Pteronotus* pectoralis muscle following glucose administration (30- and 60-min post feeding) compared to fasting (0 min). We filtered for genes with a log2 fold change > 1.5 (Fig. S1) and conducted gene set enrichment analysis using WebGestalt 2024 for Over-Representation Analysis (ORA) against KEGG, Reactome, PANTHER, and OMIM databases; results were visualized as dot plots (Fig. 1E). To examine functional organization among DEGs, we used g:Profiler to identify pathway affiliations for each gene, then clustered genes based on shared enrichment profiles using Jaccard similarity. This analysis grouped DEGs into four major functional modules: glucose utilization, lipid handling, metabolic signaling, and muscle endurance.

#### 2.2.5 Candidate Gene Expression

Expression values were normalized to the mean expression of a panel of 13 housekeeping genes (*Actb*, *Gusb*, *Hprt1*, *Ipo8*, *Ppia*, *Rpl13a*, *Rpl19*, *Rps13*, *Rps18*, *Rps23*, *Tbp*, *Tubb*, *Ywhag*), selected for their stability across all conditions (Fig. S2). This panel established a baseline reference for constitutive transcriptional activity. A curated set of candidate genes (Table S1) was selected based on their established roles in fiber-type specification, redox buffering, calcium handling, stress signaling, and muscle regeneration. Significant difference from the housekeeping mean was initially assessed using Wilcoxon signed-rank tests (Fig. S3A). To assess dynamic responses to muscle stimulation, Δlog_2_CPM values were calculated for each gene by subtracting baseline expression (WT0) from expression at WT30 and WT60. These changes were then centered relative to the mean change observed among housekeeping genes to identify candidate genes with shifts exceeding background variation. For each gene, a “z-score” was calculated using the formula:

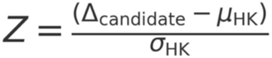

where μ and σ represent the mean and standard deviation of Δlog_2_CPM among housekeeping genes. Z-scores and two-tailed p-values were computed based on the housekeeping gene Δ distribution, followed by false discovery rate correction (Benjamini-Hochberg method). Genes with FDR < 0.05 were considered significantly responsive (Fig. S3B).

### 2.2 Isolation and Purification of bat primary myoblasts

Cryopreserved cell suspensions stored at −80° C were thawed and plated in a 24-well plate containing myoblast growth medium 1 (GM1) (Table 2). Cell cultures were maintained in a humidified incubator with 5% CO_2_ at 37° C until cell confluency reached ∼75%.

**Table 2.** Myoblasts growth media.

| Media | Base Medium | FBS | Additives | Antibiotics / Antifungals |
| --- | --- | --- | --- | --- |
| **GM1** | Ham's F-10 Nutrient Mix (Gibco, Cat# 11550043) | 20% HyClone Characterized FBS (Cytiva, Cat# SH30071.03) | 5 ng/mL recombinant human epidermal growth factor (rh EGF) (ATCC, Cat #: PCS-999-018), 10 μM dexamethasone (ATCC, Cat #: PCS-999-069), 25 μg/ml recombinant human insulin (ATCC, Cat #: PCS-999-068), 5 ng/mL recombinant human fibroblast growth factor basic protein (rh FGF-b) (ATCC, Cat #: PCS-999-020) | 2× Pen/Strep (Gibco, Cat# 15140122), 2 μg/mL Amphotericin B (R&D, Cat# B23192) |
| **GM2** | 50% Ham's F-10 Nutrient Mix +50% DMEM (ATCC, Cat# 30-2002) | 20% HyClone Characterized FBS | 5 ng/mL rh EGF, 10 μM dexamethasone, 25 μg/ml rh insulin, 5 ng/mL rh FGF-b | 2× Pen/Strep, 2 μg/mL Amphotericin B |
| **GM3** | DMEM | 20% HyClone Characterized FBS | 5 ng/mL rh EGF, 10 μM dexamethasone, 25 μg/ml rh insulin, 5 ng/mL rh FGF-b | 2× Pen/Strep, 2 μg/mL Amphotericin B |

Myoblasts were enriched using the pre-plating method, which separates cells based on differential adhesion properties to the culture surface [62–64]. Briefly, cells were seeded in myoblast growth medium on a gelatin-coated culture dish and incubated at 37° C with 5% CO_2_ for 20 min. This pre-plating step was repeated two additional times during subsequent subcultures to increase myoblast purity.

### 2.3 Cell Immortalization

Bat primary myoblasts were either spontaneously immortalized or immortalized through lentiviral overexpression of human telomerase reserve transcriptase (hTERT) and cyclin-dependent kinase 4 (CDK4). For hTERT/CDK4-mediated immortalization, primary myoblasts were transduced using lentiviral particles encoding hTERT (GenTarget Inc, Cat #: LVP1131-RB-PBS) and CDK4 (GenTarget Inc, Cat #: LVP1140-RB-PBS), each carrying a blasticidin selection marker. Transduction was performed using the spinoculation method [65]. Briefly, 50 000 bat primary myoblasts were seeded in one well of 24-well plate. The second day, primary myoblasts were spinoculated at 1000g for 2 hours with hTERT and CDK4 lentivirus, with an MOI of 20, in the presence of 8 μg/mL polybrene (Tocris Bioscience, 7711), after which, the media was replaced with fresh myoblasts media. Then, myoblasts were incubated at 37° C with 5% CO_2_. Two days post-transduction, cells were selected in 4 µg/mL blasticidin (InvivoGen, Cat #: ant-bl-1) for 6 days, at which point all the un-transduced control cells had died (Fig. S4). Both hTERT/CDK4-immortalized myoblasts (iBatM-TC1) and the spontaneously immortalized myoblasts (iBat-S1) were maintained in GM1 and subcultured at a 1:3-1:6 ratio every 2-3 days. Cells were cryobanked around every 10 passages up until passage 40 in freeze media prepared with 10% dimethyl sulfoxide (DMSO, Sigma) and 90% HyClone Characterized FBS.

### 2.4 Myotube Differentiation

For myotube differentiation, myoblasts were seeded with a cell number of 2e^5^ per well in 6-well cell culture plate (Corning) (or 1e5 per well in 24-well plate (IBIDI, Cat #: *82426*) containing 3 ml growth media and incubated as described above cell density reached >85% confluency. Then, growth media was replaced with different differentiation media (DM) to initiate differentiation (Table 3). Cell morphology changes were tracked daily following the initiation of differentiation.

**Table 3.** Myotube differentiation media.

| Media | Base Medium | Horse Serum | Additives | Antibiotics / Antifungals |
| --- | --- | --- | --- | --- |
| **DM1** | DMEM | 2% | — | 2× Pen/Strep, 2 μg/mL Amphotericin B |
| **DM2** | Ham's F-10 Nutrient Mix | 2% | — | 2× Pen/Strep, 2 μg/mL Amphotericin B |
| **DM3** | DMEM | 2% | 5 ng/mL rh EGF, 10 μM dexamethasone, 25 μg/ml rh insulin, 5 ng/mL rh FGF-b | 2× Pen/Strep, 2 μg/mL Amphotericin B |
| **DM4** | Ham's F-10 Nutrient Mix | 2% | 5 ng/mL rh EGF, 10 μM dexamethasone, 25 μg/ml rh insulin, 5 ng/mL rh FGF-b | 22× Pen/Strep, 2 μg/mL Amphotericin B |

### 2.5 Myoblasts proliferation assay

Myoblast proliferation assay was performed as previously described [29]. Briefly, myoblasts were seeded with a cell number of 1-2e^5^ per well in 6-well cell culture plate containing 3 ml different growth media: GM1, GM2 and GM3 (Table 2). Myoblasts were dissociated with TryplE and counted every day until myoblasts are >90% confluency or began differentiation.

### 2.6 Chromosome counting

Chromosome counting was performed as previously described [29]. Briefly, myoblasts were cultured in growth media in T25 cell culture flask at 37°C with 5% CO_2_, until cells density reach 60–80% confluency. Then cells were incubated in growth media containing 50 ng/ml colcemid (Invitrogen Life Technologies, Cat#: 15212–012) at 37°C with 5% CO_2_ for 2.5 hrs. Subsequently, myoblasts were washed with PBS, dissociated with TryplE, and collected through centrifuge at 300×g for 5 min. Cells were then resuspend in Hypotonic solution (1 ml growth media and 9 ml 0.075 M KCl) (Gibco, Cat#:10575-090), incubate at 37 ° C for 30 min, and centrifuged at 200×g for 5 min at RT. Eventually, the myoblasts were fixed through gradual fixation method with freshly prepared ice-cold fixative (75% Methanol, 25% glacial acetic acid). After which, cells were re-suspend in 500 µl fixative and store at 4 °C until use (usually less than 1 month). For the metaphase spreads preparation, the cell suspension was dropped onto a glass slide from a 10-cm distance and air dried. Slides were mounted with VECTASHIELD® Antifade Mounting Medium with DAPI (Vector Laboratories, Cat#: H-1200-10) and sealed with coverslip. For chromosome counting, at least 50 metaphase spreads were analyzed using fluorescence microscope. Image analysis and processing were performed using ImageJ.

### 2.7 Immunofluorescence

Cells cultured in 24-well IBIDI plate were washed with phosphate-buffered saline (PBS), fixed with 4% paraformaldehyde for 30 min at RT, washed in PBS, permeabilized for 30 min using 0.1% Triton-X, blocked for 60 min using SuperBlock™ Blocking Buffer (Thermo Scientific™, Cat #: 37580), and washed with PBST. For myoblasts marker staining, Primary PAX7 antibody (Santa Cruz, Cat #: sc-81648)(1:50) and Primary Desmin Primary (abcam, Cat #: ab15200)(1:200) was diluted in Antibody Diluent Reagent Solution (Thermo Scientific™, Cat #: 003118), added to cells, and incubated overnight at 4 °C. Cells were then washed with PBST, and incubated with CF®488A Donkey Anti-Mouse IgG (H+L)(Biotium, Cat #. 20014) and CF®647 Donkey Anti-Rabbit IgG (H+L) (Biotium, Cat #. 20047) (both diluted 1:500 in Antibody Diluent Reagent Solution) for 60 min at RT. After washing with PBST, the cell nuclei were stained with DAPI (Biolegend, Cat # 422801) (used at 1:500 dilution in PBS), for 15 min at RT, then washed with PBS. For myotube marker MF20 staining, differentiated cells were fixed, and stained as previously described, using primary MF20 antibodies (Invitrogen, Cat #: 14-6503-82) (1:200), CF®488A Donkey Anti-Mouse IgG (H+L) (Biotium, Cat #. 20014) and DAPI (Biolegend, Cat # 422801). For F-actin staining, the cells were stained with Phalloidin CF647 phalloidin conjugate (Biotium, Cat# 00041-T) (used 1:40 dilution in PBS) for 30 min at RT, then washed with PBS. Imaging was proceeded with a fluorescent microscope.

## 3. Results

### 3.1 Functional and Molecular Specializations Supporting Flight in *Pteronotus parnellii*

#### 3.1.1 Candidate gene expression reveals fiber-type heterogeneity

To investigate the structural and metabolic specializations of flight muscle, we first analyzed native pectoralis tissue using histological staining and RNA sequencing. These data provide an in vivo reference for muscle phenotype and serve as a foundation for interpreting downstream cell-based assays. Histological and transcriptomic analyses revealed a mosaic fiber-type architecture in the pectoralis muscle of *Pteronotus* (Fig. 1). Periodic acid–Schiff (PAS) staining indicated heterogeneous glycogen distribution across fibers, consistent with a mixture of slow- and fast-twitch fiber types (Fig. 1B). RNA-seq analysis confirmed co-expression of both fast-twitch and slow-twitch fiber markers (Fig. 1C). Expression of fast-twitch genes such as *Myl1, Tnnt3, Myh1* (Type IIx), *and Myh4* (Type IIb), were strongly expressed. Slow-twitch markers, including *Myl3* and *Tnnt1* were also present at relatively high levels, revealing a subset of oxidative fibers. While *Myh7* (Type I) was not detected, paralog *Myh7b* was expressed at low levels. *Myh2* (Type IIa) was not detected. Additionally, calcium-handling genes showed mixed expression: *Casq1* and *Casq2* (fast and slow paralogs of calsequestrin) were expressed at similar levels, as were *Atp2a1* (*Serca1*) and *Atp2a2* (*Serca2*) (fast and slow paralogs of sarcoplasmic reticulum Ca²⁺-ATPase). These patterns support a predominance of fast-twitch glycolytic (Type IIb) and oxidative (Type IIx) fibers, alongside hybrid calcium-handling features involving both calcium release and reuptake mechanisms typical of fast- and slow-twitch muscle.

At baseline rest (0 min), flight muscle exhibits elevated expression of candidate genes involved in redox buffering and calcium handling, as defined by levels exceeding housekeeping gene expression (Fig. S3A). These include redox-related genes *Foxo3*, *Cat*, *Sod1*, and *Txnip*, as well as calcium-handling genes noted above (*Casq1*, *Casq2*, *Atp2a1*, *Atp2a2).* While not differentially expressed across timepoints, their high expression suggests a constitutive role in supporting oxidative stability and calcium flux in muscle (Fig. 1D). *Tmc5* was included as a calcium cycling candidate gene based on its putative functional role. Its low transcript abundance suggests it may play a more limited or context-specific role in muscle physiology (Fig. S3A). These findings highlight a predominate fast-twitch fiber composition with oxidative and calcium-handling adaptations, suggesting *Pteronotus* flight muscle is tuned for both high force production and sustained performance.

#### 3.1.2 DEG analyses reveal flight muscle activation of regeneration and metabolism

To further characterize flight muscle specialization and its capacity to respond to physiological stress, we examined gene expression changes following glucose stimulation in *Pteronotus*. We focused on differentially expressed genes (DEGs) between 30 and 60 minutes post-stimulation (Fig. 1E), a window chosen to capture dynamic transcriptional responses to energy-induced stress. Using a cutoff of log_2_ fold-change >1.5 (Fig. S1), we found that most EDGs were downregulated at 60 minutes relative to 30 minutes, reflecting transient early activation of muscle development and regeneration pathways. Gene set enrichment analysis identified significant overrepresentation of terms such as muscle cell differentiation (p=5.2× 10^-5^), muscle organ development (p=7.8 × 10^-4^), and regeneration (p=0.0024), consistent with an early activation of myogenic and structural programs. Conversely, genes upregulated at 60 minutes were enriched for terms related to metabolic processes, including phosphatase complex (*p* = 6.7 × 10⁻⁴), manganese ion binding (*p* = 3.1 × 10⁻⁵), and muscle system process (*p* = 0.0012), suggesting a shift toward energy regulation and contractile function. Pathway-level enrichment analysis highlighted pathways relevant to muscle growth and glucose uptake including KEGG | PI3K-Akt signaling (*p* = 9.14 × 10⁻^3^), Panther | PI3 kinase pathway (p=0.1188), and KEGG | focal adhesion (*p* =1.83 × 10⁻^3^).

To visualize these time-dependent transitions, we generated a conceptual model overview (Fig. 1F) integrating significantly differentially expressed genes (30 and 60 min vs. baseline) and functionally relevant candidate genes with expression above housekeeping baseline (Table S1; Fig. S3). Genes were grouped into functional modules reflecting redox buffering, nutrient and stress sensing (metabolic resilience), and regeneration. For example, markers of regeneration included DEGs such as *Hey2*, *Tmc5*, *Chad, Csrp3, Lingo4, Myh4*, which peaked at 30 minutes and declined thereafter, while additional candidates with established roles in myogenic activation, cell cycle progression, and fiber-type specification *Pax7*, *Mymx*, *Etf8*, *Tmeff1*, *Maf* were significanatly upregulated compared to housekeeping levels (Fig. S3B). Genes associated with metabolic resilience included DEGs *Etv5*, *Map4k3, Slc4a7* and genes involved in redox buffering (*Nrf1*, *Pak1*, *Ucp2*) remained elevated. These dynamics suggest a coordinated transition from early muscle repair toward metabolic adaptation.

A direct comparison of 60 vs. 30 minutes post-stimulation (Fig. 1G) confirmed this temporal shift, with many regeneration-related transcripts (e.g. *Mymk, Mymx, Tmc5, and Chrng)* decreasing while metabolic regulators (e.g. *Ppm1n*, *Ppp1r3g)* increasing in expression. *Ppp1r3g*, a well-defined regulatory subunit of protein phosphatase 1, promotes glycogen synthase activation and so its upregulation facilitates glucose storage in late phase muscle activation. Notably, several highly upregulated genes at 60 minutes were uncharacterized loci (e.g., *LOC129064675*, *LOC129072723*, *LOC129068479*), potentially representing novel or bat-specific transcripts involved in late-phase muscle responses. Together, these results underscore a biphasic transcriptional response to glucose stimulation: an early-activation of regeneration and structural programs followed by a later phase of metabolic reprogramming. Given the transcriptional evidence for regenerative activation (Fig. 1E), such as elevated expression of *Pax7* and *Hey2* following glucose stimulation (Fig. S1, Fig. S3), we next sought to isolate and expand muscle progenitor cells from *P. mesoamericanus* (Fig. 2A).

**Figure 2.**
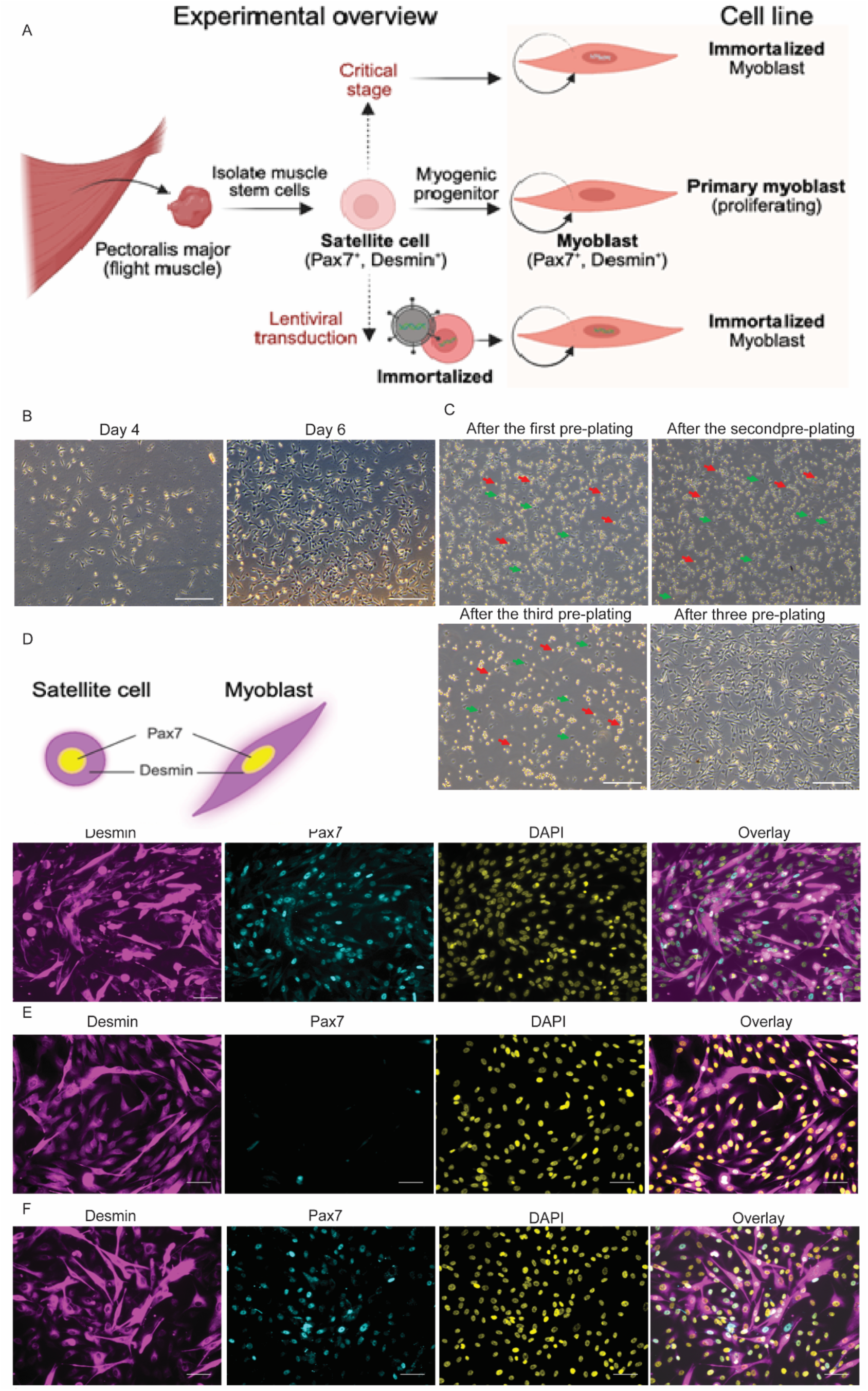
Bat primary myoblast isolation and myoblasts verification. A. Summary of the development and characterization of myoblasts cell lines. B. Primary muscle cells at 4-day and 6-day post derivation. C. Representative phase contrast images showing the muscle cells at different stages of pre-plating. Green arrow: attached cells; Red arrow: cells in suspension. D-F. Immunofluorescent staining of Desmin (magenta) and Pax7 (cyan) with DAPI counterstain (yellow) of *P. mesoamericanus* primary myoblasts, P9 (D), iBatM-Pmeso-S, P37 (E), and iBatM-Pmeso-TC, P37 (F). Scale bar: 50 μm.

### 3.2 Isolation of PAX7⁺ cells from flight muscle and verification of myogenic cells

Given the strong transcriptional evidence for regenerative activation in the skeletal muscle (Fig. 1E-G), we hypothesized that this tissue harbors a large resident population of muscle stem cells (satellite cells) that could be isolated and expanded in vitro. To test this, we adapted a pre-plating strategy commonly used for myoblast enrichment in mammalian systems [66–69]. Using pre-plating, we derived stable, self-renewing satellite cells from the pectoralis major muscle of *P. mesoamericanus*.

Primary cells began attaching to the cell culture dish 3-4 days after seeding (Fig. 2B). By day 6, cultures reached 80% confluence with mix of spindle- and triangle-shaped cells (Fig. 2B). After 3 rounds of pre-plating, the adherent cell population became uniform in morphology (Fig. 2C), typical of activated satellite cells and early myoblasts, suggesting successful enrichment of myogenic progenitors. Immunofluorescence staining confirmed co-expression of muscle stem cell marker PAX7 and the intermiedate filament protein Desmin, validating the presence of activated muscle stem cells (Fig. 2D). In addition, when switched to growing in standard differentiation media (DM1), these cells readily fused into multinucleated myotubes within 48 hours (Fig. 3A). Myotube identity was confirmed by immunostaining for myosin heavy chain (MyHC), a myotube marker; Fig. 3B, 3C). Interestingly, many differentiated myotubes exhibited frequent spontaneous contraction as early as two days post-differentiation (Video 1), indicating functional integrity and neuromuscular activity.

**Figure 3.**
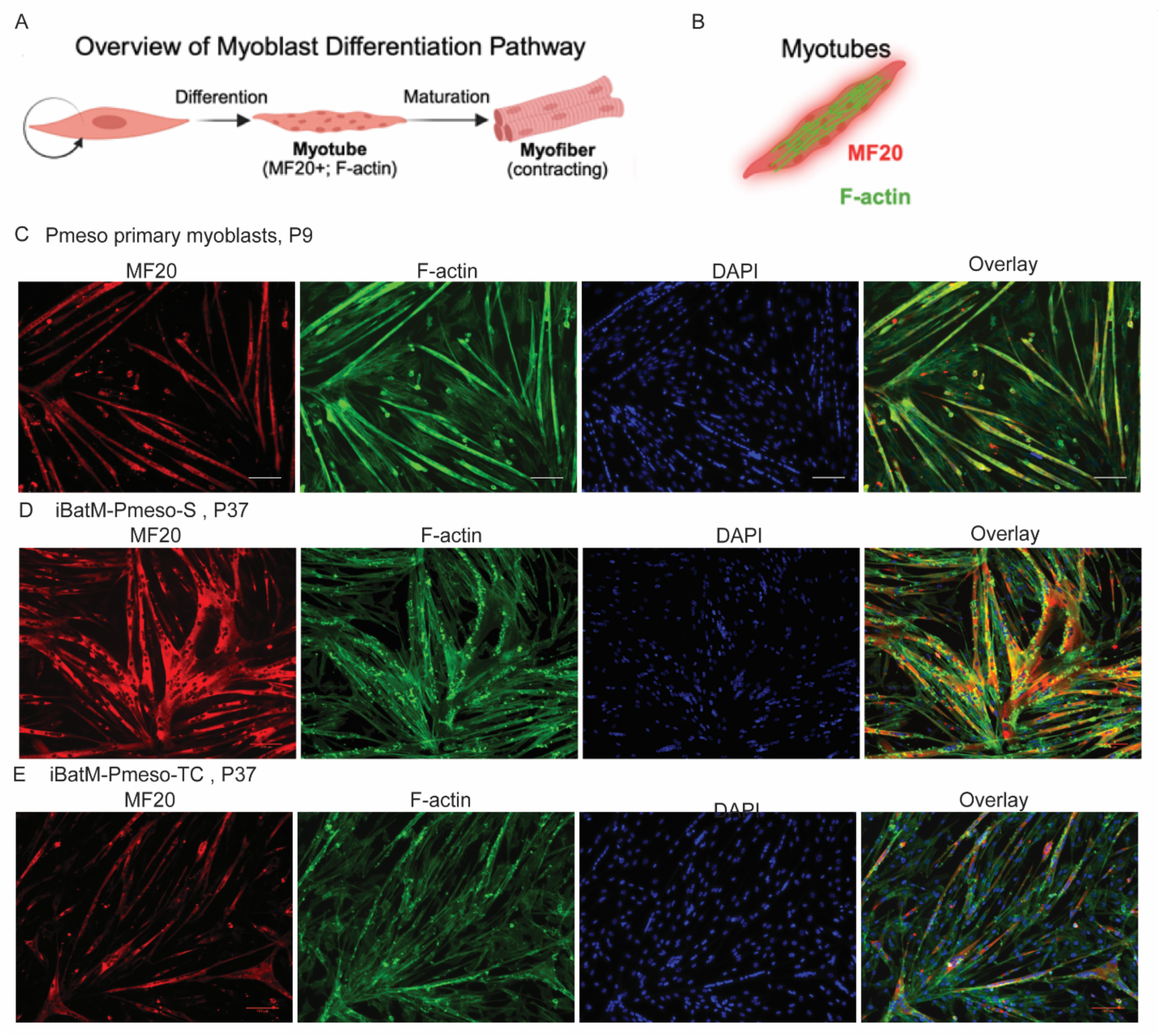
Bat Myoblast Lines Retain Differentiation Capacity. A. Schematic overview of myoblast differentiation into multinucleated myotubes and subsequent maturation into myofibers. B. Expression of a myogenic signature marker in differentiated myotubes, shown by immunostaining for myosin heavy chain (MF20; red) and counterstaining with F-actin (green).C–E. Immunofluorescent staining of myosin heavy chain (MF20; red), F-actin (green), and nuclei (DAPI; blue) in differentiated Pmeso primary myoblasts, P9 (C), iBatM-Pmeso-S, P37 (D), and iBatM-Pmeso-TC, P37 (E), demonstrating preserved myogenic potential across cell types. Scale bar: 100 μm.

### 3.3 Establishment of immortalized bat myoblast (iBatM) cell lines

Following successful isolation and validation of primary myogenic progenitors, we next aimed to establish long-term expandable muscle cell lines. To establish immortalized bat cell lines, primary *P. mesoamericanus* myoblasts were co-transduced with lentiviral constructs expressing human telomerase reverse transcriptase (hTERT) and cylcin-dependent kinase 4 (CDK4), each containing a blasticidin selection marker. The resulting engineered line, iBatM-Pmeso-TC, showed stable proliferation over multiple passages (Fig. S4A).

In parallel, a separate population of non-transduced primary cells entered a transient proliferative crisis before recovering and resuming growth, giving rise to a spontaneously immortalization line, iBatM-Pmeso-S. Both lines have been maintained for over 50 passages across 5 months in culture, with no observable changes in cellular morphology (Fig. S4B). These results confirm successful immortalization by both engineered and spontaneous mechanisms.

To assess whether these lines retained myogenic identity, we performed immunofluorescence staining for the canonical markers PAX7 and Desmin. The hTERT/CDK4-immortalized iBatM-Pmeso-TC line co-expressed both markers at passage 37 (Fig. 2F), consistent with an activated satellite cell state, similar to primary cell cultures (Fig. 2D). In contrast, the spontaneous line iBatM-Pmeso-S retained Desmin expression with low or absent PAX7 population (Fig. 2E), indicating a more committed myoblast phenotype with reduced stemness. These results demonstrate that the two immortalized bat cell lines can be derived through two mechanisms and that they capture distinct stages of the myogenic lineage, providing flexible and renewable in vitro models for muscle biology and regeneration.

### 3.4 Immortalized bat myoblasts are genetically stable

To validate genomic stability of the immortalized lines, we performed karyotype analysis on both primary and immortalized long-term passaged cells. Primary *P. mesoamericanus* myoblasts at passage 9 (P9) exhibited a diploid chromosome number of 38 (2N = 38), consistent with our previous karyotyping of bat fibroblasts [29] and published karyotypes for *Pteronotus* species (Fig. 4A). Both immortalized cell lines, the engineered iBatM-Pmeso-TC and the spontaneous iBatM-Pmeso-S, retained the same diploid count at passage 37 (P37) (Fig. 4B-C), indicating that extended passaging did not compromise chromosomal integrity. These results confirm that the immortalized bat myoblasts remain genetically stable over time, supporting their use as a reliable in vitro platform.

**Figure 4.**
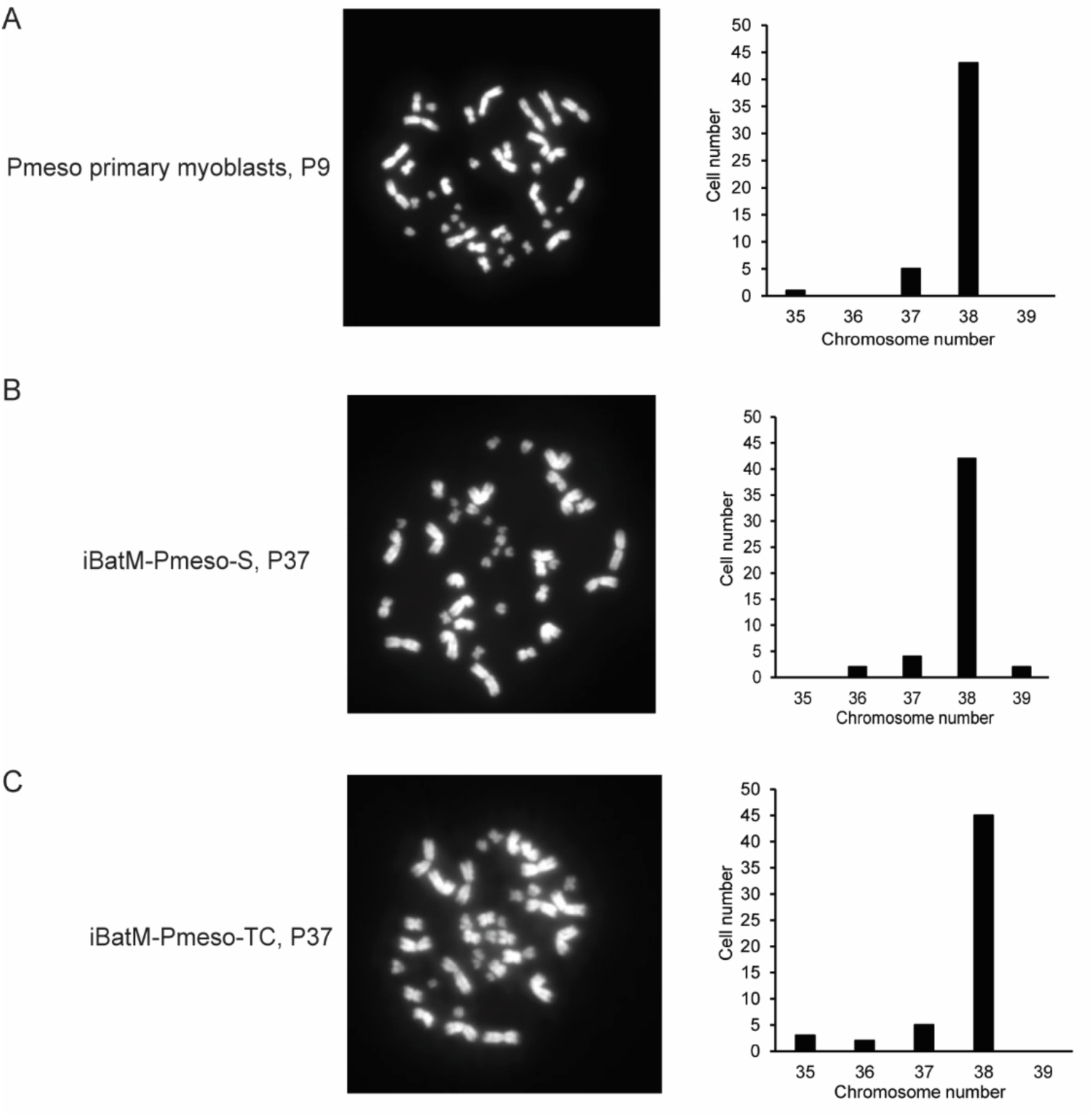
Bat Myoblast Lines Retain Genetic Stability After Extended Passaging. A. Representative chromosome spread and count distribution for *P. mesoamericanus* primary myoblasts (Pmeso) at passage 9. A total of 49 metaphase spreads were analyzed, with a diploid karyotype of 2n=38. **B**. Representative chromosome spread and counts for iBatM-Pmeso-S at late passage (P37). Of 50 metaphase spreads, most maintained the expected diploid number (2n=38), indicating genomic stability despite extended culture. **C.** Representative image of chromosome spread and chromosome counts for iBatM-Pmeso-TC at P37. Among 55 metaphase spreads, the majority maintained 2n=38, consistent with long-term karyotypic stability.

### 3.5 Immortalized bat myoblasts retain the proliferation capacity of bat primary myoblasts

To evaluate the long-term proliferative stability of the immortalized lines, we measured doubling time at early (P8), mid (P20), and late (P40) passages. Both cell iBatM-Pmeso-TC and iBatM-Pmeso-S proliferated well in standard F10-based growth medium (GM1) across all timepoints (Fig. 5, Fig. S5A, B). At P8, doubling times were 23.42 hrs for iBatM-Pmeso-TC and 26.33 hrs for iBatM-Pmeso-S (Fig. 5A, 5B). At P20, both cell lines showed slower proliferation: 31.07 hrs for iBatM-Pmeso-TC and 37.22 hrs for iBatM-Pmeso-S, following by recovery at P40 (25.46 hrs and 28.96 hrs, respectively; Fig. 5A, 5B). These trends indicate that both cell lines maintain stable growth profiles after the initial adaptation to culture, with rates comparable to primary bat myoblasts.

**Figure 5.**
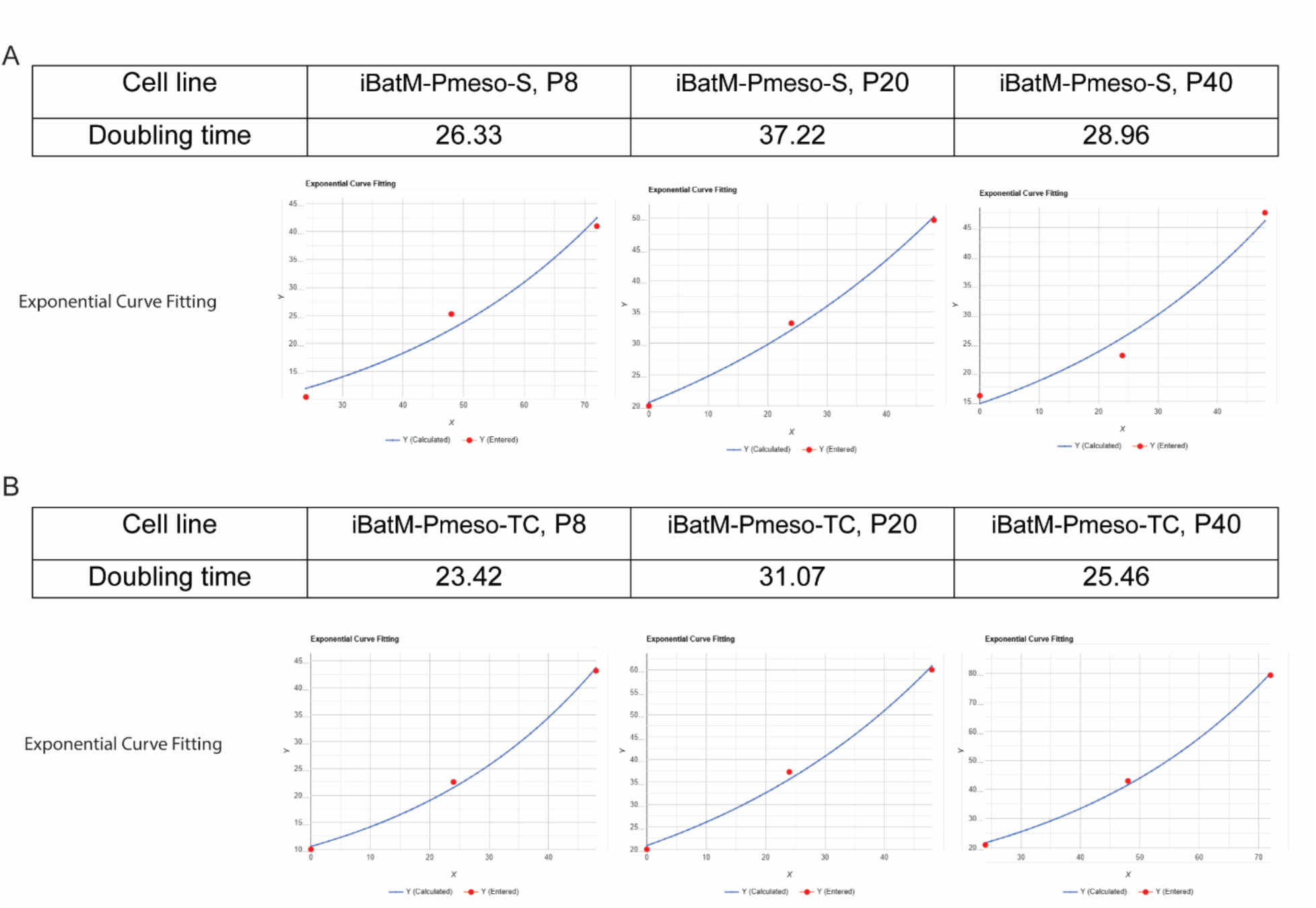
Spontaneous and hTERT/CDK4 Immortalization Support Long-Term Proliferation. A. Doubling time and exponential curve fitting for spontaneously immortalized myoblasts iBatM-Pmeso-S at passages 8 (P8), P20, and P40, demonstrating the bypass of senescence. B. Doubling time and exponential growth curve fitting for hTERT/CDK4-immortalized myoblasts iBatM-Pmeso-TC at P8, P20, and P40, showing stable proliferation. Doubling Time Computing software: https://www.doubling-time.com/compute_more.php

We next tests how basal media composition affects cell proliferation and differention potential. Previous studies have shown that different growth media can influence myoblasts behavior, including stemness maintenance and initiation of differentiation [63, 64, 71]. To systematically assess this in bat myoblasts, both cell lines were cultured in three growth media formulations: F10 (GM1), DMEM/F10 (1:1; GM2), or DMEM (GM3), each were supplemented with identical growth factors and serum (Table 2). Morphological assessment revealed no notable differences across conditions (Fig. S5A, B), but cell counts showed that both cell lines proliferated more rapidly in GM2 and GM3 compared to GM1 (Fig. S5C).

In addition, cells in GM2 and GM3 began differentiating into myotubes once cultures reached >95% confluence, with widespread myotube formation observed by day 3 and detachment day 8 (Fig. S6, Fig. S7). In contrast, cells maintained in GM1 remained largely undifferentiated even at day 8 under high-density conditions, showing minimal myotube formation. These results suggest that bat myoblasts lines retain long-term proliferative capacity and respond to media compoistion in a predictable manner, growing faster and initiating differentiation in DMEM-based media, while maintaing a more undifferentiated, stem-like state in F10-based condidtions.

### 3.6 Immortalized bat Myoblasts Retain the Differentiation Capacity of bat Primary Myoblasts

To evaluate whether the immortalized lines retained their capacity to differentiate into functional muscle fibers, iBatM-Pmeso-TC and iBatM-Pmeso-S were cultured in standard differentiation media (DM1). Both cell lines successfully formed multinucleated myotubes within two days (Fig. S8, Fig. S9) and spontaneously myotube contractions were observed by day 3 (Video 2 and Video 3). Immunofluorescence staining confirmed expression of MyHC, validating their myogenic identity (Fig. 3D, E). Notably, the spontaneously immortalized iBatM-Pmeso-S line formed significantly more myotubes under the same conditions, suggesting differences in differentiation efficiency between the two lines.

To further examine how culture conditions influence differentiation, we systematically tested four differentiation media formulations (DM1-DM4; Table 3) that varied in basal media (DMEM vs F10). Both cell lines formed mature multi-nucleated myotubes in DMEM-based media (DM1 and DM3), but not in F10-based media (DM2 and DM4), even with extended culture (Fig. S8, Fig. S9). This finding is consistent with our previous observation that F10 supports a more stem-like, undifferentiated state, while DMEM promotes differentiation.

Since myoblasts fuse into myotubes in GM3, we further evaluated the effects of serum (2% horse serum versus 20% HyClone Characterized FBS) on bat myotube differentiation. . Cells cultured in GM3, DM1, and DM3 formed functional myotubes, while no myotube formation was seen in GM1, DM2 and DM4, (Fig. S8B-C, Fig. S9B-C). This suggests that serum reduction is not required for differentiation in these bat myoblast lines, and that basal media composition is the primer driver differentiation competence. Together, these findings demonstrate that both immortalized myoblast lines retain functional differentiation capacity and contractile potential, and that differentiation efficiency is strongly influenced by basal media composition.

### 3.7 Immortalized Bat Myoblasts Enable Functional Profiling of Flight Muscle Under Metabolic Overload

The establishment of spontaneously contracting myotubes from both primary (Video 1) and immortalized cells (Video 2 and 3) offers a unique in vitro system to study how bat skeletal muscle supports the extreme physiological demands of powered flight. While detailed mechanistic assays are ongoing, the robust and early-onset contractile activity observed during differentiation suggests enhanced functional maturation, potentially involving neuromuscular junction (NMJ) priming or sustained excitation–contraction coupling. Such spontaneous contraction is rarely observed under standard culture conditions (Table 4), underscoring the distinctive physiology of bat myotubes.

**Table 4.**
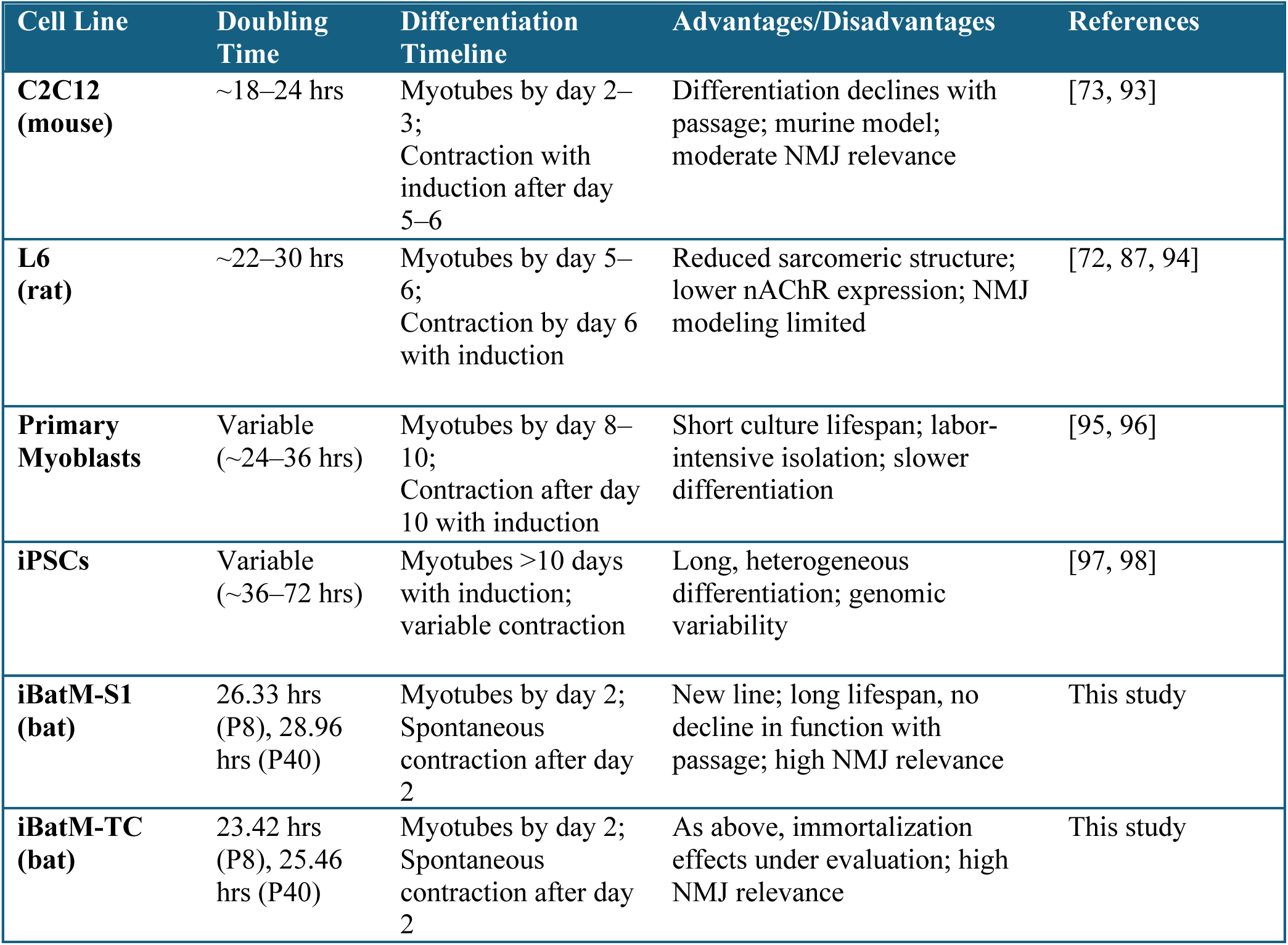
Comparative features of commonly used skeletal muscle cell models.

To contextualize this cellular behavior in a physiological context, we examined whole-animal metabolic response to glucose overload in *P. mesoamericanus* using the standardized glucose tolerance tests (GTT), a non-lethal assay of systemic metabolic function. Following glucose challenge, *P. mesoamericanus* maintained tightly regulated blood glucose levels, indicating robust systemic glucose (Fig. 6). Because lethal sampling was not permitted in *P. mesoamericanus*, we turned to *P. parnellii*, a closely related species that shares similar dietary ecology and key metabolic features of nutrient assimilation (Camacho and Bernal 2024). Biochemical profiling revealed that flight muscle energy reserves were primarily stored as triglycerides, with relatively lower levels of glycogen (Fig. 6A), consistent with a lipid-based endurance metabolism. In contrast to P. mesoamericanus, GTT for *P. parnellii* showed broader inter-individual variation and frequent sustained hyperglycemia (Fig. 6B). Using a linear mixed-effects model, we confirmed significant species-level differences in glucose clearance (*p* < 0.001).

**Figure 6.**
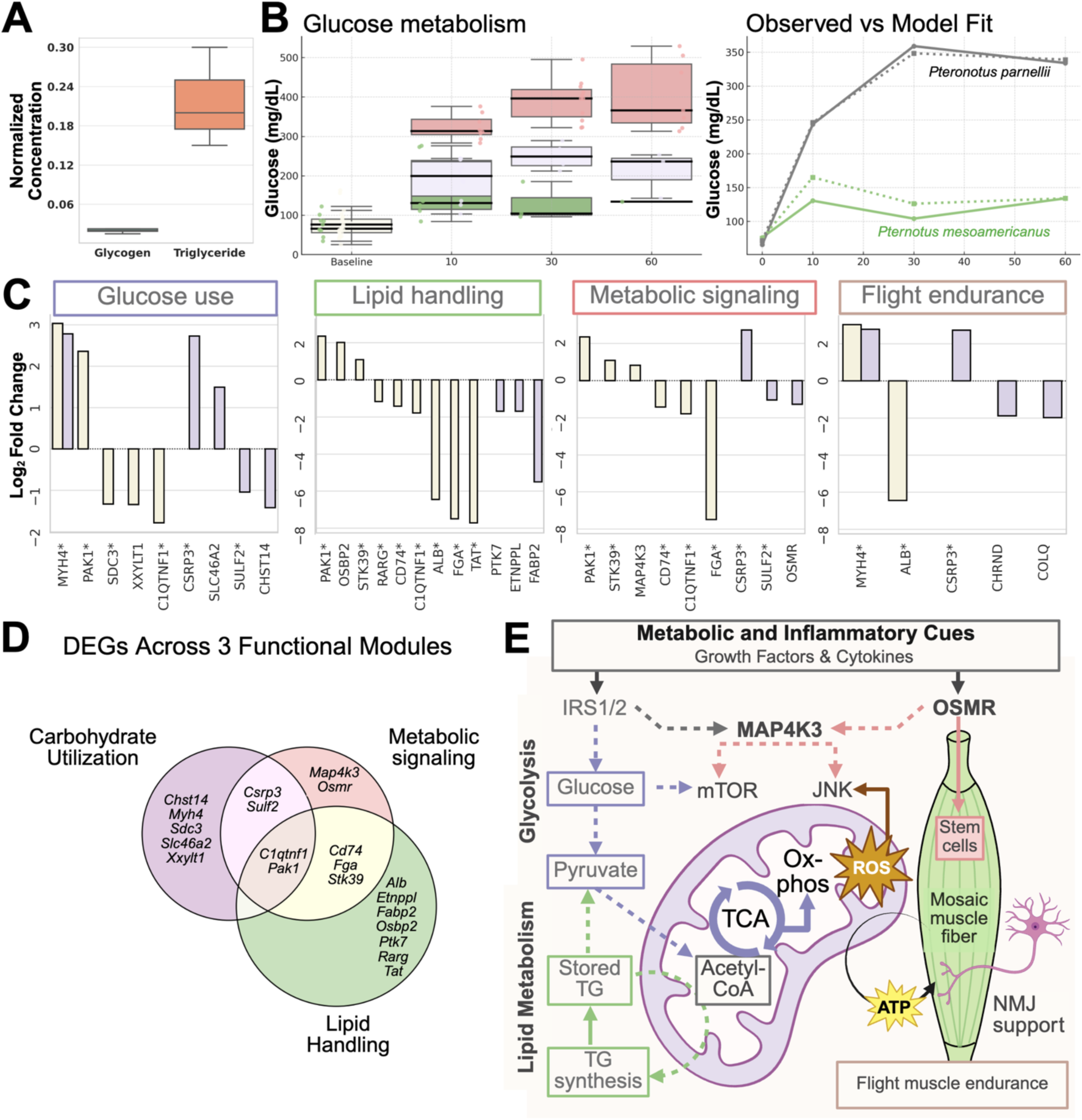
Integrated molecular, physiological, and functional adaptations in flight muscle. A. Normalized concentrations of glycogen and triglycerides measured in *P. parnellii* pectoralis muscle, showing preferential storage of triglycerides. B. Glucose tolerance test (GTT) time series. Left: Blood glucose levels measured at baseline, 10, 30, and 60 minutes after glucose administration across individual bats (n=36). *P. mesoamericanus* (green) shows a uniformly low response, while *P. parnellii* exhibits both high and low glucose clearance phenotypes. Right: Linear mixed-model fit shows a significant species × time interaction (*p* < 0.001) in glucose clearance for *P. mesoamericanus* (green, n=10) and *Pteronotus parnellii* (gray, n=26). C. Log_2_ fold change in gene expression for differentially expressed genes (DEGs) in flight muscle at 30 min (yellow) or 60 min (purple) post-glucose administration compared to baseline (0 min). Only DEGs with |log_2_FC| > 1.5 and adjusted p-value < 0.01 (EdgeR) are shown. DEGs are grouped into four functional modules: Glucose Use, Lipid Handling, Metabolic Signaling, and Flight Endurance based on enrichment clustering of g:Profiler-enriched terms. Genes with (*) are shared in different modules. Module-specific DEGs include *Chst14, Slc46a2, Xxylt1* (glucose handling) and *Osbp2*, *Ptk7*, *Etnppl*, *Fabp2* (lipid handling), highlighting the transcriptional basis of metabolic specialization in bat flight muscle. D. Venn diagram illustrating overlap of DEGs across three functional modules: Carbohydrate Utilization, Lipid Handling, and Metabolic Signaling. Genes in overlapping regions are implicated in multiple pathways. Notably, *Map4k3* and *Osmr* are uniquely assigned, representing upstream metabolic and cytokine signaling inputs. E. Schematic of core cellular metabolic pathways integrating transcriptomic data into a functional framework for flight muscle performance. This conceptual model incorporates well-established features of glucose metabolism, lipid handling, mitochondrial ATP production, and neuromuscular activity, based on general principles of cell biology. Growth factor and cytokine signals (*IRS1/2* and *OSMR)* activate downstream *MAP4K3*, all supported at the transcript level. These signaling pathways converge on *mTOR* and *JNK* to regulate metabolic and stress responses. Pyruvate-derived acetyl-CoA is routed to the TCA cycle or triglyceride (TG) synthesis. Stored TG serves as an energy reserve for mitochondrial energy production. Lipid handling DEGs *Fabp2* and *Etnppl* support mitochondrial beta-oxidation and mitochondrial membrane integrity during TG mobilization. ATP production supports neuromuscular junction (NMJ) activity and contraction, with postsynaptic genes *Chrnd* and *Colq* (endurance-associated) and contractile genes *Myh4* and *Csrp3* (contraction mediated glucose uptake) upregulated. OSMR also contributes to regenerative signaling, consistent with stem cell activation.

To identify underlying molecular programs, we returned to transcriptomic analysis of *P. parnellii* pectoralis muscle between 30 and 60 minutes post-glucose stimulation. Differential expression analysis (|log_2_FC| > 1.5, FDR < 0.01) revealed genes associated with fuel utilization and stress response (Fig. S1). Functional clustering of enriched pathways, based on Jaccard similarity among g:Profiler-enriched terms, grouped DEGs into four major functional modules: glucose utilization, lipid handling, metabolic signaling, and flight endurance (Fig. 6C). Key lipid handling genes such as *Fabp2* and *Etnppl* suggest active beta oxidation and mitochondrial membrane integrity during triglyceride mobilization. Endurance related genes *Chrnd* and *Colq*, as well as contractile genes *Myh4* and *Csrp3,* support contraction-coupled glucose uptake.

A Venn diagram illustrating gene assignments across modules showed substantial overlap (Fig. 6D), underscoring the coordinated nature of muscle responses to nutrient load. Notably, signaling regulators *Map4k3* (nutrient sensing) and *Osmr* (cytokine-mediated) were uniquely assigned, pointing to upstream metabolic and regenerative processes. To further contextualize molecular findings, we developed a schematic model of core cellular pathways supporting flight muscle performance (Fig. 6E). Upstream metabolic and cytokine signaling pathways act through *IRS1/2* and *OSMR*, both of which were transcriptionally upregulated following glucose stimulation. These signals converge on nutrient sensitive regulators such as *MAP4K3* and downstream effectors mTOR and JNK, which mediate stress and metabolic responses. Within this postprandial state, pyruvate-derived acetyl-CoA is directed into the TCA cycle or diverted toward triglyceride (TG) synthesis, aligning with our biochemical data showing TG-enriched flight muscle.

Lipid-driven ATP production supports ongoing contraction and neuromuscular junction (NMJ) activity, as reflected in the enriched expression of postsynaptic genes (*Chrnd*, *Colq*) and contractile regulators (*Myh4*, *Csrp3*). In parallel, *OSMR* contributes to regenerative signaling through activation of muscle stem cell pathways, consistent with our earlier evidence of Pax7⁺ progenitor activation. By synthesizing cellular phenotypes, physiological assays, and dynamic transcriptional responses, this multi-scale framework highlights how bat flight muscle coordinates metabolic flexibility, contractile function, and stem cell activity to sustain muscle performance.

## 4 Discussion

Skeletal muscle is a metabolically dynamic tissue essential for locomotion, energy homeostasis, and repair. In flying mammals, these demands are intensified by the energetic and mechanical load of powered flight. To investigate how bat muscle accommodates such stress, we first performed transcriptomic profiling of *Pteronotus* pectoralis muscle before and after glucose stimulation—a physiological proxy for flight energy overload (Fig. 1). This in vivo analysis revealed enriched expression of genes involved in calcium handling (e.g., *Casq1, Atp2a1*), redox buffering (*Foxo3, Cat, Sod1*), membrane transport (*Slc16a1*), and nutrient sensing (*Igf1, Nr1d1*), as well as neuromuscular components (*Chrnd, Colq*) and contractile regulators (*Myh4, Csrp3*). These findings define molecular programs that support energy production, oxidative resilience, and sustained contraction under metabolic load, laying the foundation for understanding muscle performance in the context of flight. To enable direct experimental access to these mechanisms, we next established the first immortalized bat myoblast cell lines. These lines serve as a renewable in vitro platform to study muscle development, regeneration, and metabolism in a species naturally adapted for powered flight.

Myoblasts, the progenitor cells of skeletal muscle, are essential for muscle development, injury repair, and regeneration [2, 3]. In the present study, we report the first successful establishment of bat myoblast cell lines. These bat lines retain hallmarks of myogenic identity, including PAX7 and Desmin expression, and readily differentiate into multinucleated myotubes. Remarkably, the differentiated myotubes exhibited spontaneous contractions in vitro—suggesting intrinsic properties of excitability and contractile readiness that may reflect specialized flight muscle physiology. Isolating primary myoblasts from pectoralis muscle is technically challenging due to the mix of myoblasts, fibroblasts, and other differentiated cell types. Fibroblasts and myoblasts are both proliferative and share morphology, which complicates enrichment, particularly in non-model species lacking validated markers and reagents. We used a pre-plating method based on differential adhesion to enrich myoblasts, following protocols established in other systems [66–69].

We established two immortalized bat myoblast cell lines: iBatM-Pmeso-S through spontaneous immortalization and iBatM-Pmeso-TC via hTERT/CDK4 overexpression. These strategies are informed by prior models (Table 1) such as L6 cells, C2C12 [73], hTERT-immortalized myoblasts from human [36, 37, 46, 47] and canine sources [49], and spontaneous immortalization of fish myoblasts [51, 74, 75]. Both bat cell lines retain myogenic potential and stable karyotypes, with doubling times of ∼25-29 hours at passage 40. Notably, iBatM-Pmeso-TC preserved a higher proportion of PAX7+ progenitor cells than iBatM-Pmeso-S or even primary myoblasts, suggesting that hTERT/CDK4 overexpression may enhance stemness by bypassing senescence pathways. Having two immortalized cell lines that differ in progenitor maintenance provides a valuable framework for studying myogenic states and their regulation under various metabolic or physiological stress.

We optimized cell culture media to balance proliferation and differentiation based on insight from other systems [51, 76, 77]. Growth media supplemented with fibroblast growth factor (FGF), epidermal growth factor (EGF), insulin, and dexamethasone media might promote robust proliferation and stemness preservation. Among basal media tested, F10-based media was most effective at maintaining undifferentiated progenitor states, while DMEM promoted myotube differentiation. The contrast is likely driven by DMEM’s higher concentrations of glucose (4.5 g/L vs. 1.1 g/L in F-10), as glucose restriction below 0.9 g/L has been shown to block myotube formation [78] [79]. DMEM also contains calcium and essential amino acids (leucine, valine, and lysine), factors known to promote mTOR signaling and myoblast fusion [80–82]. In contrast to DMEM media, F10 instead contains a broader range of micronutrients and amino acids that support stemness, including copper, zinc, hypoxanthine, lipoic acid, alanine, asparagine, aspartic acid, glutamic acid, proline, biotin, and vitamin B12 [83, 84]. These findings emphasize the importance of species-specific formulations and suggest that F10 media may be well suited for regenerative studies, while DMEM is more appropriate for differentiation protocols.

Importantly, we found base media composition had a stronger effect on differentiation than serum concentration. While low serum conditions are typically used to induce myotube differentiation in vitro [37, 51, 71, 85, 86], our results show that myoblasts cultured in F10-based media failed to differentiate even under low serum condition. In contrast, cells in DMEM or DMEM/F10 rapidly formed functional myotubes regardless of serum levels. This suggests that nutritional cues (e.g. glucose and amino acids) may be a primary driver of myotube fusion in bat myoblasts, with implications for interpreting metabolic signaling during bat muscle development.

The transcriptomic profiling observed in native *Pteronotus* pectoralis tissue closely align with our in vitro findings. Upregulated genes involved in calcium handling (*Casq1*, *Casq2*, *Atp2a1*, *Atp2a2*), redox buffering (*Foxo3*, *Cat*, *Sod1*), membrane transport (*Slc16a1*), and nutrient sensing (*Igf1*, *Nr1d1*), and contractile function (*Myh4*, *Csrp3*) mirror the functional phenotype of spontaneous myotube contraction. A systems-level model (Fig. 6E) integrates these signals, highlighting how upstream nutrient and cytokine pathways (e.g. insulin via IRS1/2, OSM vis OSMR) converge on downstream effectors such as MAP4K3, mTOR, and JNK.

Taken together, these transcriptomic and functional data validate the physiological relevance of our immortalized cell lines and provide the foundation for future mechanistic studies for flight muscle biology. These data lay the foundation for downstream experimental studies—including metabolic flux (assays of glucose and fatty acid oxidation), mitochondrial dynamics, and insulin-independent glucose uptake—to directly test hypotheses about metabolic specialization in flying mammals. More broadly, the availability of these renewable lines now enables a unique opportunity to investigate lineage-specific muscle adaptations in a clade of mammals that remain largely unexplored in vertebrate muscle biology.

Finally, the establishment of these bat myoblast lines provides a new experimental system alongside standard rodent lines such as C2C12 and L6 (Table 4). While rodent models have been invaluable for studying muscle differentiation and insulin signaling, they originate from short-lived terrestrial, glycolytic mammals with limited oxidative capacity. In contrast, *P. mesoamericanus* is a flight-adapted species characterized by high muscular endurance, physiological resilience, and longevity. The stable bat cell lines enables comparative studies to investigate conserved and lineage-specific muscle adaptations, with potential relevance to aging, metabolic disease, and therapeutic muscle repair.

## 5 Conclusions

In this study, we isolated primary myoblasts from wild-caught bat pectoralis muscle and successfully established two genetically stable myoblasts cell lines. This is the first foundational in vitro platform to study muscle regeneration and function in a flying mammal. Both immortalized bat cell lines retain proliferative capacity and readily differentiate into functional myotubes, exhibiting spontaneous contraction. Comparative optimization of culture conditions revealed that DMEM supports myoblasts proliferation and myotube differentiation, while F10 media more effectively preserves the progenitor state, offering flexibility for applications focused on muscle repair vs maintenance. Finally, by integrating in vitro cell behavior, in vivo physiology, and transcriptomic profiling, we uncovered a biphasic muscle response to metabolic overload in bats—characterized by early regenerative activation, sustained contractile readiness, and a reliance on triglyceride-based energy stores. By capturing these coordinated responses across scales, the iBatM cell lines offer a powerful system to investigate both conserved and lineage-specific strategies that support muscle performance under extreme physiological stress and evolutionary adapataions for powered flight.

## Author Contributions

Fengyan Deng: Data curation, formal analysis, methodology, validation, visualization, manuscript writing and revision. Valentina Peña: field investigation, metabolic assays, and RNA-seq; Pedro Morales-Sosa: field study and RNA-seq; Andrea Bernal-Rivera: GTT field study; Bowen Yang: RNA-seq data analysis. Shengping Huang, Sonia Ghosh, Maria Katt, Luciana Castellano: Data curation and manuscript editing. Cindy Maddera: visualization; Zulin Yu: formal analysis. Nicolas Rohner: Funding acquisition and supervision. Chongbei Zhao: Conceptualization, investigation, supervision, manuscript writing and revision. Jasmin Camacho: Conceptualization, funding acquisition, project administration, supervision, field study, metabolic assays, data analysis, manuscript writing and revision.

## Funding

Field work and glucose tolerance tests (GTT) were funded by the National Science Foundation (NSF) grant no. 2109717 (2021–2022) awarded to JC. RNA-sequencing was funded by the Burroughs Wellcome Fund (BWF) Postdoctoral Diversity Enrichment Program No. G-1022339 (2022-2025) to JC. Metabolic assays and additional research resources were provided by the Howard Hughes Medical Institute (HHMI) Hanna H. Gray Fellows Program no. GT15991 (2022-2025) to JC. Additional support for AB-R, PM-S, and VP was provided by National Institutes of Health (NIH) grant no. 1DP2AG071466-0 to NR. VP was supported by the Graduate School *Biology Scholars Program* at the Stowers Institute during this work (2023-2024). Any opinions, findings, and conclusions or recommendations expressed in this material are those of the authors and do not necessarily reflect the views of the NIH, NSF, BWF, or HHMI.

## Institutional Review Board Statement

Not applicable.

## Acknowledgments

The fieldwork was carried out within the framework of the informally dubbed “Belize Bat-a-thon”, founded by Dr. Brock Fenton and co-organized by Dr. Nancy Simmons and we thank everyone for the field support. The authors would like to thank the Stowers Institute for Medical Research technology centers: Cells, Tissues, and Organoids Center members for the meaningful discussion throughout protocol optimization; We also thank Dai Tsuchiya, Seth Malloy and Kexi Yi for the meaningful discussion on the in vitro model characterization.

## Data Availability Statement

Original data underlying this manuscript can be accessed from the Stowers Original Data Repository at https://www.stowers.org/research/publications/LIBPB-2563.

## Conflicts of Interest

The authors declare no conflicts of interest to declare.

## Abbreviations

hTERT: human telomerase reverse transcriptase
CDK4: cyclin-dependent kinase 4
CDKI: cyclin-dependent kinase inhibitor
rh EGF: recombinant human epidermal growth factor
rh FGF-b: recombinant human fibroblast growth factor basic protein
P/S: Penicillin/Streptomycin
P. meso: Pteronotus mesoamericanus

---

**Table S1:** A curated list of candidate genes included in the illustrated model of flight muscle at rest and following glucose stimulation (Fig. 1), with functional annotations and known roles in skeletal muscle.

| Gene | Functional Module | Relative to HK | DEG | Biological Role in Muscle |
| --- | --- | --- | --- | --- |
| **Atp2a1** | Calcium cycling | > HK | No | Calcium reuptake in fast-twitch muscle |
| **Atp2a2** | Calcium cycling | > HK | No | Calcium handling in slow/oxidative muscle |
| **Casq1** | Calcium cycling | > HK | No | Calcium buffering in sarcoplasmic reticulum (fast muscle) |
| **Casq2** | Calcium cycling | > HK | No | Calcium buffering (cardiac/oxidative muscle) |
| **Cat** | Redox | > HK | No | ROS detoxification enzyme (catalase) |
| **Foxo3** | Redox | > HK | Yes | Redox regulation, muscle atrophy signaling |
| **Sod1** | Redox | > HK | No | ROS scavenging (superoxide dismutase) |
| **Txnip** | Redox | > HK | No | Redox regulation and glucose sensing |
| **Tmc5** | Muscle regeneration | < HK | Yes | Transmembrane channel; unclear function in muscle |
| **Mymx** | Muscle regeneration | > HK | Yes | Myoblast fusion |
| **Myh4** | Calcium cycling | > HK | Yes | Fast-twitch muscle fiber myosin |
| **Pak1** | Energy sensing | > HK | Yes | Energy sensing, insulin signaling |
| **Plk2** | Stress sensing | > HK | Yes | Stress-responsive kinase |
| **Chad** | Muscle regeneration | > HK | Yes | ECM remodeling in skeletal muscle |
| **Osbp2** | Lipid handling | > HK | Yes | Oxysterol binding, lipid sensing |
| **Csrp3** | Muscle regeneration | > HK | Yes | Z-disk protein, structural integrity |
| **Lingo4** | Muscle regeneration | > HK | Yes | Poorly characterized; possibly involved in regeneration |
| **Hey2** | Muscle regeneration | > HK | Yes | Myogenic transcription factor (Notch target) |
| **Pax7** | Muscle regeneration | > HK | Yes | Muscle stem cell maintenance |
| **Ucp2** | Redox | > HK | Yes | Mitochondrial uncoupling, redox regulation |
| **Tmeff1** | Muscle regeneration | > HK | Yes | Muscle regeneration and differentiation |
| **Map4k3** | Energy sensing | > HK | Yes | Nutrient signaling (mTOR, JNK) |
| **Nrf1** | Redox | > HK | Yes | Mitochondrial biogenesis and oxidative metabolism |
| **Slc4a7** | Energy sensing | > HK | Yes | Proton-coupled ion transporter (pH regulation) |
| **Maf** | Redox | > HK | Yes | Redox-responsive transcription factor |
| **Prkaa2** | Energy sensing | > HK | Yes | AMPK catalytic subunit (energy sensing) |
| **Etv5** | Stress sensing | < HK | Yes | Stress-responsive transcription factor |
| **Sdc3** | Muscle regeneration | < HK | Yes | Cell adhesion, muscle stem cell regulation |
| **Entpd3** | Stress sensing | < HK | Yes | Purinergic signaling; immune/stress response |

Video 1. Contraction of P2 *P. Meso* myotubes from primary myoblasts.

Video 2. Contraction of P43 self-immortalized (iBatM-Pmeso-S) differentiated myotube.

Video 3. Contraction of P43 hTERT/CDK4-immortalized (iBatM-Pmeso-TC) differentiated myotube.

## Notes

### Competing Interest Statement

The authors have declared no competing interest.

## References

1. Chal, J. and O. Pourquie, Making muscle: skeletal myogenesis in vivo and in vitro. Development, 2017. 144(12): p. 2104–2122.

2. Joe, A.W., et al., Muscle injury activates resident fibro/adipogenic progenitors that facilitate myogenesis. Nat Cell Biol, 2010. 12(2): p. 153–63.

3. Wosczyna, M.N., et al., Mesenchymal Stromal Cells Are Required for Regeneration and Homeostatic Maintenance of Skeletal Muscle. Cell Rep, 2019. 27(7): p. 2029–2035 e5.

4. Giannini, N.P., et al., Palaeoatmosphere facilitates a gliding transition to powered flight in the Eocene bat, Onychonycteris finneyi. Commun Biol, 2024. 7(1): p. 365.

5. Matthew F. Jones, K.C.B.N.B.S., Phylogeny and systematics of early Paleogene bats. Journal of Mammalian Evolution, 2024. 31(2): p. 18.

6. Rummel, A.D., S.M. Swartz, and R.L. Marsh, Thermal Stability of Contractile Proteins in Bat Wing Muscles Explains Differences in Temperature Dependence of Whole-Muscle Shortening Velocity. Physiol Biochem Zool, 2023. 96(2): p. 100–105.

7. Sears, K.E., et al., Development of bat flight: morphologic and molecular evolution of bat wing digits. Proc Natl Acad Sci U S A, 2006. 103(17): p. 6581–6.

8. Cretekos, C.J., et al., Regulatory divergence modifies limb length between mammals. Genes Dev, 2008. 22(2): p. 141–51.

9. Tokita, M., T. Abe, and K. Suzuki, The developmental basis of bat wing muscle. Nat Commun, 2012. 3: p. 1302.

10. Eckalbar, W.L., et al., Transcriptomic and epigenomic characterization of the developing bat wing. Nat Genet, 2016. 48(5): p. 528–36.

11. Feigin, C.Y., et al., Convergent deployment of ancestral functions during the evolution of mammalian flight membranes. Sci Adv, 2023. 9(12): p. eade7511.

12. Shen, Y.Y., et al., Adaptive evolution of energy metabolism genes and the origin of flight in bats. Proc Natl Acad Sci U S A, 2010. 107(19): p. 8666–71.

13. Zhang, G., et al., Comparative analysis of bat genomes provides insight into the evolution of flight and immunity. Science, 2013. 339(6118): p. 456-60.

14. Cao, T. and J.P. Jin, Evolution of Flight Muscle Contractility and Energetic Efficiency. Front Physiol, 2020. 11: p. 1038.

15. Cooper, L.N., et al., Bats as instructive animal models for studying longevity and aging. Ann N Y Acad Sci, 2024. 1541(1): p. 10–23.

16. Johnson, A.A., M.N. Shokhirev, and B. Shoshitaishvili, Revamping the evolutionary theories of aging. Ageing Res Rev, 2019. 55: p. 100947.

17. Hua, R., et al., Experimental evidence for cancer resistance in a bat species. Nat Commun, 2024. 15(1): p. 1401.

18. Athar, F., et al., Limited cell-autonomous anticancer mechanisms in long-lived bats. Nat Commun, 2025. 16(1): p. 4125.

19. Ahn, M., et al., Unique Loss of the PYHIN Gene Family in Bats Amongst Mammals: Implications for Inflammasome Sensing. Sci Rep, 2016. 6: p. 21722.

20. Ahn, M., et al., Bat ASC2 suppresses inflammasomes and ameliorates inflammatory diseases. Cell, 2023. 186(10): p. 2144–2159 e22.

21. Moreno Santillan, D.D., et al., Large-scale genome sampling reveals unique immunity and metabolic adaptations in bats. Mol Ecol, 2021. 30(23): p. 6449–6467.

22. Morales, A.E., et al., Bat genomes illuminate adaptations to viral tolerance and disease resistance. Nature, 2025. 638(8050): p. 449-458.

23. Faure, P.A., D.E. Re, and E.L. Clare, Wound Healing in the Flight Membranes of Big Brown Bats Journal of Mammalogy, 2009. 90(90): p. 1148–1156.

24. Ma, S., et al., Cell culture-based profiling across mammals reveals DNA repair and metabolism as determinants of species longevity. Elife, 2016. 5.

25. Stawski, C., C.K.R. Willis, and G. F., The importance of temporal heterothermy in bats. Journal of Zoology, 2014. 292(2): p. 86–100.

26. Bernal-Rivera, A., Cuellar-Valencia, O.M., Calvache-Sánchez, C. and Murillo-García, O.E.,, Morphological, anatomical, and physiological signs of senescence in the great fruit-eating bat (Artibeus lituratus). Acta Chiropterologica, 2002. 24(2): p. 405–413.

27. Irving, A.T., et al., Optimizing dissection, sample collection and cell isolation protocols for frugivorous bats. Methods in Ecology and Evolution, 2020. 11(1): p. 150–158.

28. Carvalho, V.S., et al., Isolation and establishment of skin-derived and mesenchymal cells from south American bat Artibeus planirostris (Chiroptera - Phyllostomidae). Tissue Cell, 2021. 71: p. 101507.

29. Deng, F., et al., Establishing Primary and Stable Cell Lines from Frozen Wing Biopsies for Cellular, Physiological, and Genetic Studies in Bats. Curr Protoc, 2024. 4(9): p. e1123.

30. Jagannathan, N.S., et al., Multi-omic analysis of bat versus human fibroblasts reveals altered central metabolism. Elife, 2024. 13.

31. Alcock, D., et al., Generating bat primary and immortalised cell-lines from wing biopsies. Sci Rep, 2024. 14(1): p. 27633.

32. Gonzalez, V., et al., Expanding the bat toolbox: Carollia perspicillata bat cell lines and reagents enable the characterization of viral susceptibility and innate immune responses. PLoS Biol, 2025. 23(4): p. e3003098.

33. Rozwadowska, N., et al., Optimization of human myoblasts culture under different media conditions for application in the in vitro studies. Am J Stem Cells, 2022. 11(1): p. 1–11.

34. Ding, S., et al., Characterization and isolation of highly purified porcine satellite cells. Cell Death Discov, 2017. 3: p. 17003.

35. Shahini, A., et al., Efficient and high yield isolation of myoblasts from skeletal muscle. Stem Cell Res, 2018. 30: p. 122–129.

36. Lathuiliere, A., et al., Immortalized human myoblast cell lines for the delivery of therapeutic proteins using encapsulated cell technology. Mol Ther Methods Clin Dev, 2022. 26: p. 441–458.

37. Mamchaoui, K., et al., Immortalized pathological human myoblasts: towards a universal tool for the study of neuromuscular disorders. Skelet Muscle, 2011. 1: p. 34.

38. Harley, C.B., A.B. Futcher, and C.W. Greider, Telomeres shorten during ageing of human fibroblasts. Nature, 1990. 345(6274): p. 458-60.

39. Renault, V., et al., Regenerative potential of human skeletal muscle during aging. Aging Cell, 2002. 1(2): p. 132–9.

40. Shammas, M.A., Telomeres, lifestyle, cancer, and aging. Curr Opin Clin Nutr Metab Care, 2011. 14(1): p. 28–34.

41. Teixeira, M.Z., Telomere length: biological marker of cellular vitality, aging, and health-disease process. Rev Assoc Med Bras (1992), 2021. 67(2): p. 173–177.

42. Bodnar, A.G., et al., Extension of life-span by introduction of telomerase into normal human cells. Science, 1998. 279(5349): p. 349-52.

43. Vaziri, H. and S. Benchimol, Reconstitution of telomerase activity in normal human cells leads to elongation of telomeres and extended replicative life span. Curr Biol, 1998. 8(5): p. 279–82.

44. Rayess, H., M.B. Wang, and E.S. Srivatsan, Cellular senescence and tumor suppressor gene p16. Int J Cancer, 2012. 130(8): p. 1715–25.

45. Zhu, C.H., et al., Cellular senescence in human myoblasts is overcome by human telomerase reverse transcriptase and cyclin-dependent kinase 4: consequences in aging muscle and therapeutic strategies for muscular dystrophies. Aging Cell, 2007. 6(4): p. 515–23.

46. Stadler, G., et al., Establishment of clonal myogenic cell lines from severely affected dystrophic muscles - CDK4 maintains the myogenic population. Skelet Muscle, 2011. 1(1): p. 12.

47. Nunez-Manchon, J., et al., Immortalized human myotonic dystrophy type 1 muscle cell lines to address patient heterogeneity. iScience, 2024. 27(6): p. 109930.

48. Robin, J.D., et al., Isolation and immortalization of patient-derived cell lines from muscle biopsy for disease modeling. J Vis Exp, 2015(95): p. 52307.

49. Tone, Y., et al., Immortalized Canine Dystrophic Myoblast Cell Lines for Development of Peptide-Conjugated Splice-Switching Oligonucleotides. Nucleic Acid Ther, 2021. 31(2): p. 172–181.

50. Saad, M.K., et al., Continuous fish muscle cell line with capacity for myogenic and adipogenic-like phenotypes. Sci Rep, 2023. 13(1): p. 5098.

51. Xue, T., et al., A spontaneously immortalized muscle stem cell line (EfMS) from brown-marbled grouper for cell-cultured fish meat production. Commun Biol, 2024. 7(1): p. 1697.

52. Clare, E.L., et al., Diversification and reproductive isolation: cryptic species in the only New World high-duty cycle bat, Pteronotus parnellii. BMC Evol Biol, 2013. 13: p. 26.

53. Riskin, D.K., et al., Quantifying the complexity of bat wing kinematics. J Theor Biol, 2008. 254(3): p. 604–15.

54. Ospina-Garcés, S.M., et al., The relationship between wing morphology and foraging guilds: exploring the evolution of wing ecomorphs in bats. Biological Journal of the Linnean Society, 2023. 142(4): p. 481–498.

55. Smotherman, M. and A. Guillen-Servent, Doppler-shift compensation behavior by Wagner’s mustached bat, Pteronotus personatus. J Acoust Soc Am, 2008. 123(6): p. 4331–9.

56. Schnitzler, H.U. and A. Denzinger, Auditory fovea and Doppler shift compensation: adaptations for flutter detection in echolocating bats using CF-FM signals. J Comp Physiol A Neuroethol Sens Neural Behav Physiol, 2011. 197(5): p. 541–59.

57. Chionh, Y.T., et al., High basal heat-shock protein expression in bats confers resistance to cellular heat/oxidative stress. Cell Stress Chaperones, 2019. 24(4): p. 835–849.

58. Scheben, A., et al., Long-Read Sequencing Reveals Rapid Evolution of Immunity- and Cancer-Related Genes in Bats. Genome Biol Evol, 2023. 15(9).

59. Camacho, J., et al., Sugar assimilation underlying dietary evolution of Neotropical bats. Nat Ecol Evol, 2024. 8(9): p. 1735–1750.

60. Elizarraras, J.M., et al., WebGestalt 2024: faster gene set analysis and new support for metabolomics and multi-omics. Nucleic Acids Res, 2024. 52(W1): p. W415-W421.

61. Kolberg, L., et al., *g:Profiler-interoperable web service for functional enrichment analysis and gene identifier mapping (2023 update)*. Nucleic Acids Res, 2023. 51(W1): p. W207–W212.

62. Stephens, D.C., et al., Protocol for isolating mice skeletal muscle myoblasts and myotubes via differential antibody validation. STAR Protoc, 2023. 4(4): p. 102591.

63. Zygmunt, K., et al., Influence of Media Composition on the Level of Bovine Satellite Cell Proliferation. Animals (Basel), 2023. 13(11).

64. Neyroud, N., et al., Gene delivery to cardiac muscle. Methods Enzymol, 2002. 346: p. 323–34.

65. Anastasov, N., et al., Optimized Lentiviral Transduction Protocols by Use of a Poloxamer Enhancer, Spinoculation, and scFv-Antibody Fusions to VSV-G. Methods Mol Biol, 2016. 1448: p. 49–61.

66. Yoshioka, K., et al., A Modified Pre-plating Method for High-Yield and High-Purity Muscle Stem Cell Isolation From Human/Mouse Skeletal Muscle Tissues. Front Cell Dev Biol, 2020. 8: p. 793.

67. Kim, S.H., et al., Optimal Pre-Plating Method of Chicken Satellite Cells for Cultured Meat Production. Food Sci Anim Resour, 2022. 42(6): p. 942–952.

68. Qu, Z., et al., Development of approaches to improve cell survival in myoblast transfer therapy. J Cell Biol, 1998. 142(5): p. 1257–67.

69. Gharaibeh, B., et al., Isolation of a slowly adhering cell fraction containing stem cells from murine skeletal muscle by the preplate technique. Nat Protoc, 2008. 3(9): p. 1501–9.

70. Simmons, N.B.a.C., T.M., Phylogenetic Relationships Of Mormoopid Bats (Chiroptera: Mormoopidae) Based On Morphological Data. Bulletin of the American Museum of Natural History, 2001. 2001(258): p. 1–100.

71. Sh., Z., et al., Characterization of Proliferation Medium and Its Effect on Differentiation of Muscle Satellite Cells from Larimichthys crocea in Cultured Fish Meat Production. Fishes, 2023. 8(9).

72. Yaje, D., Retention of differentiation potentialities during prolonged cultivation of myogenic cells. Proc Natl Acad Sci U S A, 1968. 61(2): p. 477–83.

73. Yaje, D. and O. Saxel, Serial passaging and differentiation of myogenic cells isolated from dystrophic mouse muscle. Nature, 1977. 270(5639): p. 725-7.

74. Long, X., et al., Establishment and characterization of a skeletal myoblast cell line of grass carp (Ctenopharyngodon idellus). Fish Physiol Biochem, 2023. 49(5): p. 1043–1061.

75. Krishnan, S., et al., Establishment and Characterization of Continuous Satellite Muscle Cells from Olive Flounder (Paralichthys olivaceus): Isolation, Culture Conditions, and Myogenic Protein Expression. Cells, 2023. 12(18).

76. Doumit, M.E., D.R. Cook, and R.A. Merkel, Fibroblast growth factor, epidermal growth factor, insulin-like growth factors, and platelet-derived growth factor-BB stimulate proliferation of clonally derived porcine myogenic satellite cells. J Cell Physiol, 1993. 157(2): p. 326–32.

77. Milasincic, D.J., et al., Stimulation of C2C12 myoblast growth by basic fibroblast growth factor and insulin-like growth factor 1 can occur via mitogen-activated protein kinase-dependent and -independent pathways. Mol Cell Biol, 1996. 16(11): p. 5964–73.

78. Dugdale, H.F., et al., The role of resveratrol on skeletal muscle cell differentiation and myotube hypertrophy during glucose restriction. Mol Cell Biochem, 2018. 444(1-2): p. 109–123.

79. Fulco, M., et al., Glucose restriction inhibits skeletal myoblast differentiation by activating SIRT1 through AMPK-mediated regulation of Nampt. Dev Cell, 2008. 14(5): p. 661–73.

80. Duan, Y., et al., Effect of branched-chain amino acid ratio on the proliferation, differentiation, and expression levels of key regulators involved in protein metabolism of myocytes. Nutrition, 2017. 36: p. 8–16.

81. Jin, C.L., et al., mTORC1 Mediates Lysine-Induced Satellite Cell Activation to Promote Skeletal Muscle Growth. Cells, 2019. 8(12).

82. Sinha, S., Y. Elbaz-Alon, and O. Avinoam, Ca(2+) as a coordinator of skeletal muscle differentiation, fusion and contraction. FEBS J, 2022. 289(21): p. 6531–6542.

83. Ichio, 2nd, I. Kimura, and E. Ozawa, Promotion of Myoblast Proliferation by Hypoxanthine and RNA in Chick Embryo Extract: hypoxanthine/RNA/myoblast proliferation/embryo extract). Dev Growth Dijer, 1985. 27(2): p. 101–110.

84. Ohashi, K., et al., Zinc promotes proliferation and activation of myogenic cells via the PI3K/Akt and ERK signaling cascade. Exp Cell Res, 2015. 333(2): p. 228–237.

85. Duran, B., M. Dal-Pai-Silva, and D. Garcia de la Serrana, Rainbow trout slow myoblast cell culture as a model to study slow skeletal muscle, and the characterization of mir-133 and mir-499 families as a case study. J Exp Biol, 2020. 223(Pt 2).

86. Luo, Y., et al., Simplifying the protocol for low-pollution-risk, efficient mouse myoblast isolation and differentiation. Adv Biotechnol (Singap), 2025. 3(1): p. 8.

87. Richler, C. and D. Yaje, The in vitro cultivation and differentiation capacities of myogenic cell lines. Dev Biol, 1970. 23(1): p. 1–22.

88. Lopez, S.M., et al., Creation and characterization of an immortalized canine myoblast cell line: Myok9. Mamm Genome, 2020. 31(3-4): p. 95–109.

89. Baik, J., et al., Establishment of Skeletal Myogenic Progenitors from Non-Human Primate Induced Pluripotent Stem Cells. Cells, 2023. 12(8).

90. Ikeda, D., Y. Otsuka, and N. Kan-No, Development of a novel Japanese eel myoblast cell line for application in cultured meat production. Biochem Biophys Res Commun, 2024. 734: p. 150784.

91. Guo, D., et al., Establishment and Characterization of a Chicken Myoblast Cell Line. Int J Mol Sci, 2024. 25(15).

92. Perujo, A., et al., Establishment and characterization of the Cuvier’s beaked whale (Ziphius cavirostris) myogenic cell line. Res Vet Sci, 2025. 182: p. 105471.

93. Blau, H.M., C.P. Chiu, and C. Webster, Cytoplasmic activation of human nuclear genes in stable heterocaryons. Cell, 1983. 32(4): p. 1171–80.

94. Charest, A., B.H. Wainer, and P.R. Albert, Cloning and differentiation-induced expression of a murine serotonin1A receptor in a septal cell line. J Neurosci, 1993. 13(12): p. 5164–71.

95. Rando, T.A. and H.M. Blau, Primary mouse myoblast purification, characterization, and transplantation for cell-mediated gene therapy. J Cell Biol, 1994. 125(6): p. 1275–87.

96. Bentzinger, C.F., et al., Cellular dynamics in the muscle satellite cell niche. EMBO Rep, 2013. 14(12): p. 1062–72.

97. Majioletti, S.M., et al., Efficient derivation and inducible differentiation of expandable skeletal myogenic cells from human ES and patient-specific iPS cells. Nat Protoc, 2015. 10(7): p. 941–58.

98. Chal, J., et al., Generation of human muscle fibers and satellite-like cells from human pluripotent stem cells in vitro. Nat Protoc, 2016. 11(10): p. 1833–50.

