## Extended Data for "From Development to Regeneration: Insights into Flight Muscle Adaptations from Bat Muscle Cell Lines"

*Chongbei Zhao

Stowers Institute for Medical Research, Kansas City, MO, USA

* Jasmin Camacho

Stowers Institute for Medical Research, Kansas City, MO, USA

### Supplemental Figures and Legends

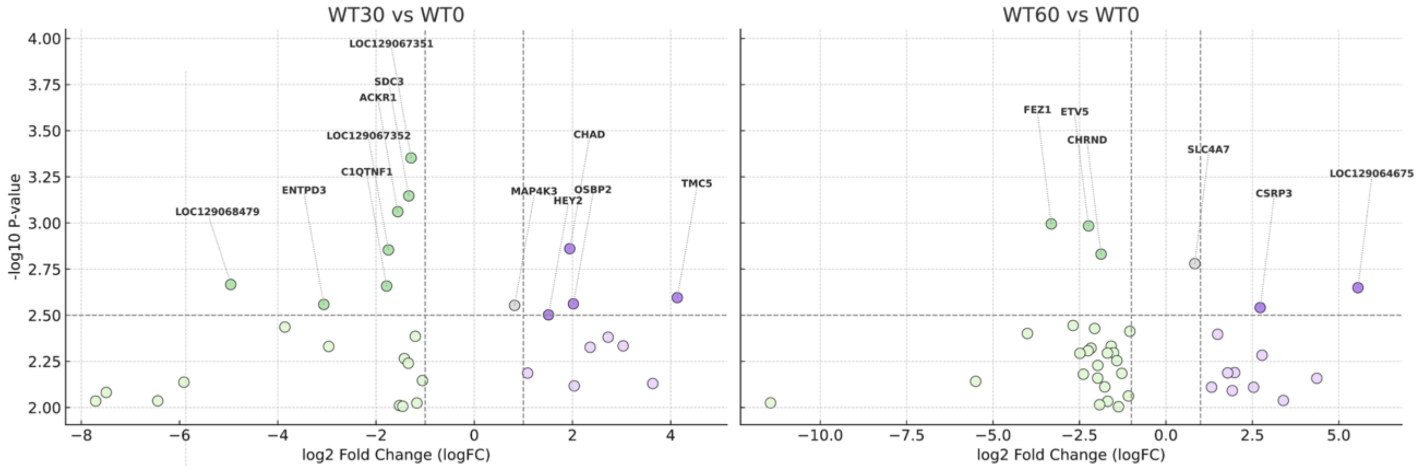


**Figure S1: Differential gene expression in bat flight muscle following glucose stimulation.** Volcano plots showing differential gene expression between 30 minutes post-glucose (WT30) and baseline (WT0) and between 60 minutes post-glucose (WT60) and baseline (WT0). Each point represents a gene. Genes with absolute $\log_2$ fold change > 1 are colored by statistical significance: darker violet (upregulated) and green (downregulated) indicate $-\log_{10}$(p-value) > 2.5 ($p < 0.0032$); lighter shades indicate the same fold change with lower statistical support ($-\log_{10}$(p-value) ≤ 2.5). Genes with $|\log_2$ fold change$| \leq 1$ are shown in gray. Dashed lines denote fold change thresholds (±1) and the significance cutoff ($p = 0.0032$). Genes passing both thresholds are labeled. These include transcription factors, signaling molecules, and membrane-associated proteins hypothesized to contribute to metabolic remodeling and stress responses in flight muscle. Left: Upregulated genes include *Chad*, *Sdc3*, *Heye2*, *Map4k3*, *Osbp2*, and *Tmc5*, associated with membrane remodeling, regeneration, and intracellular signaling. Downregulated genes include *LOC129067351* and *LOC129067352*, which show homology to MHC class II histocompatibility antigens (DR and DQ chains), suggesting transient suppression of immune activation. *Etv5*, a stress-responsive transcription factor, also decreases. Right: Upregulated genes include *Slc4a7* and *Csrp3*, linked to ion homeostasis and cytoskeletal organization. *LOC124900675*, a strongly upregulated uncharacterized non-coding RNA, is noted as a candidate regulatory transcript.

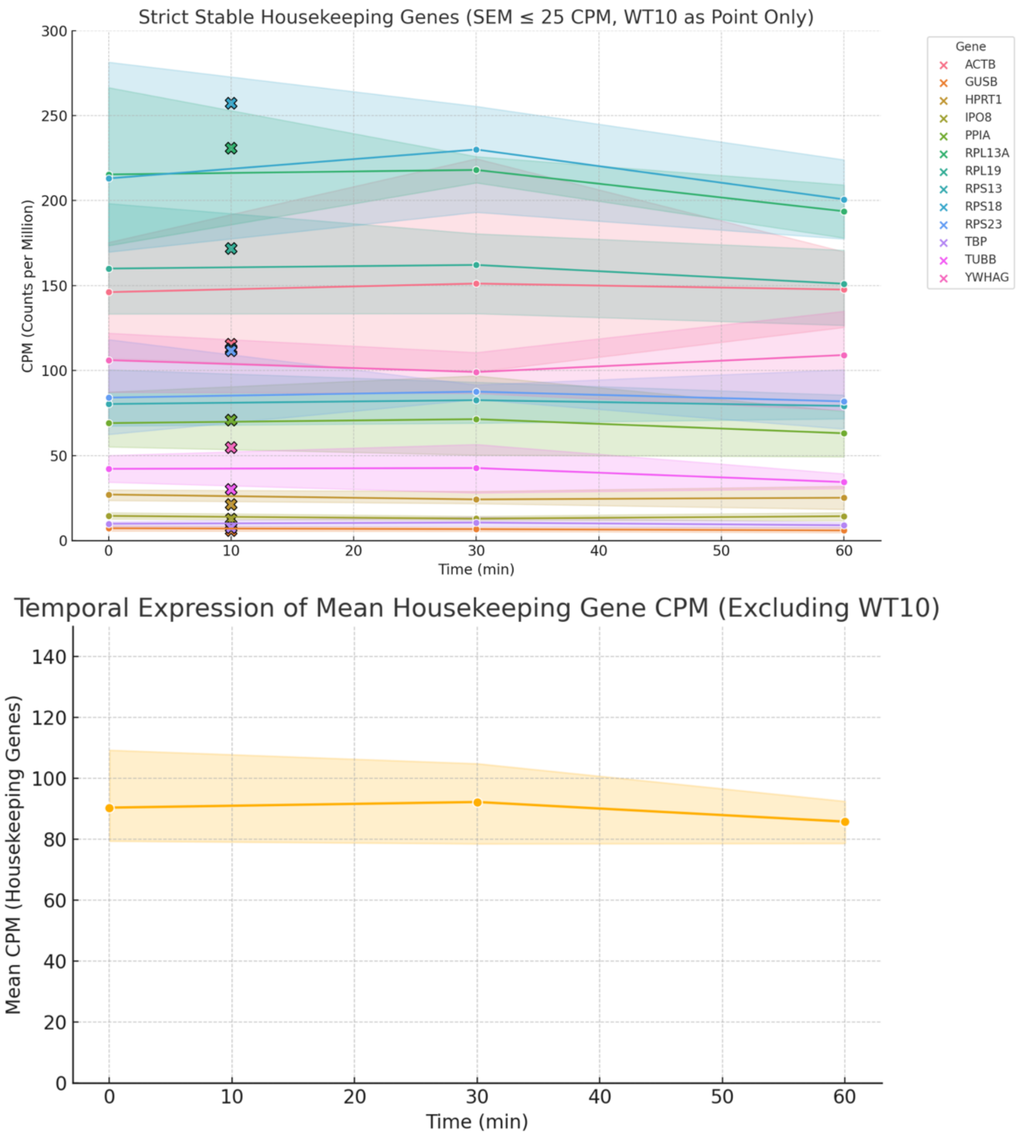


**Figure S2. Housekeeping genes are stable across conditions. (Top)** Counts per million (CPM) for individual housekeeping genes (n = 13) across timepoints 0, 30, and 60 minutes post-stimulation. Genes were selected based on expression stability (SEM ≤ 25 CPM) and average expression ≥ 25 CPM. Shaded regions represent SEM. WT10 values are shown as points only and were excluded from model fitting due to limited replication. **(Bottom)** Mean CPM for all strict housekeeping genes (excluding WT10), showing stable expression across time. Shading denotes SEM. These genes serve as a reference for identifying biologically meaningful expression above baseline transcriptional activity.

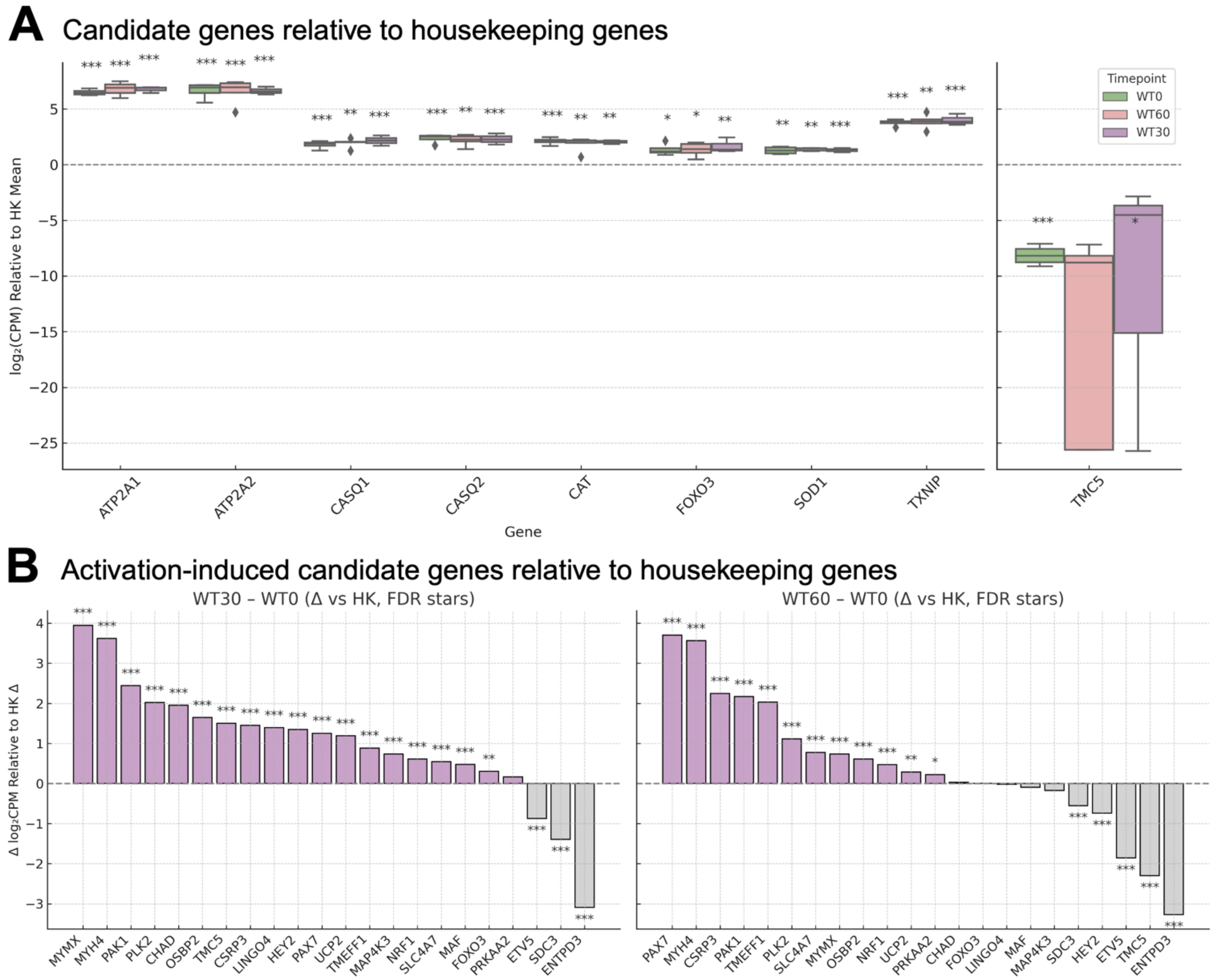


**Figure S3: Gene expression changes relative to housekeeping genes.** A. Boxplots show $\log_2$-transformed CPM (counts per million) values of candidate genes involved in redox buffering (*Foxo3, Cat, Sod1, Txnip*) and calcium cycling (*Casq1, Casq2, Atp2a1, Atp2a2, Tmc5*) across three timepoints (WT0, WT30, WT60). Values are centered relative to the mean of 13 housekeeping genes (HK = 0 reference line). A dashed line at 0 reflects this HK baseline, and colors denote timepoint (green = WT0, purple = WT30, pink = WT60). Asterisks indicate statistical significance relative to HK expression at each timepoint (*FDR < 0.05, **FDR < 0.01, ***FDR < 0.001; two-sided t-test with Benjamini-Hochberg correction). *Tmc5* is shown in a separate panel to accommodate its broader expression range. B. Bar plots show the activation-induced gene expression changes with $\Delta\log_2$CPM for each DEG at WT30 and WT60, relative to the average expression change observed in housekeeping genes. Bars are colored based on directionality (purple: upregulated; gray: downregulated). A dashed line at 0 indicates no change relative to housekeeping gene baseline dynamics. Asterisks denote significance based on FDR-adjusted p-values (*FDR < 0.05, **FDR < 0.01, ***FDR < 0.001). Genes are sorted by relative expression change magnitude at each timepoint.

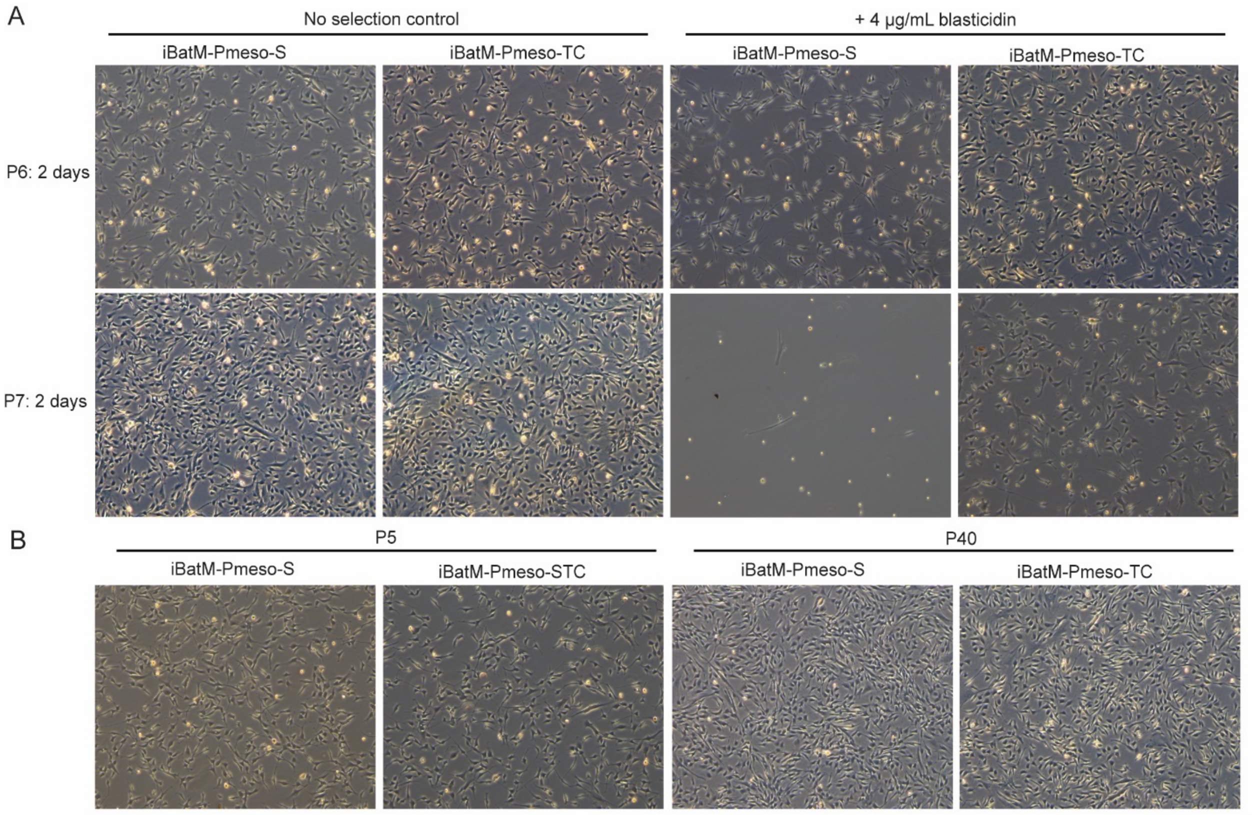


**Figure S4. Bat myoblast immortalization**. A. Phase-contrast images of primary myblasts from *P. mesoamericanus (*iBatM-Pmeso-S) and hTERT/CDK4 lentivirus transduced myoblasts (iBatM-Pmeso-TC) treated with or without 4 µg/mL blasticidin. Images were taken 2 days post-treatment across multiple passages (P6-7) to assess drug sensitivity and cell viability. B. Phase-contrast images showing the morphology of iBatM-Pmeso-S, and iBatM-Pmeso-TC early (passage 5, left) and late (passage 40, right) stages. Scale bar: 100 µm.

A

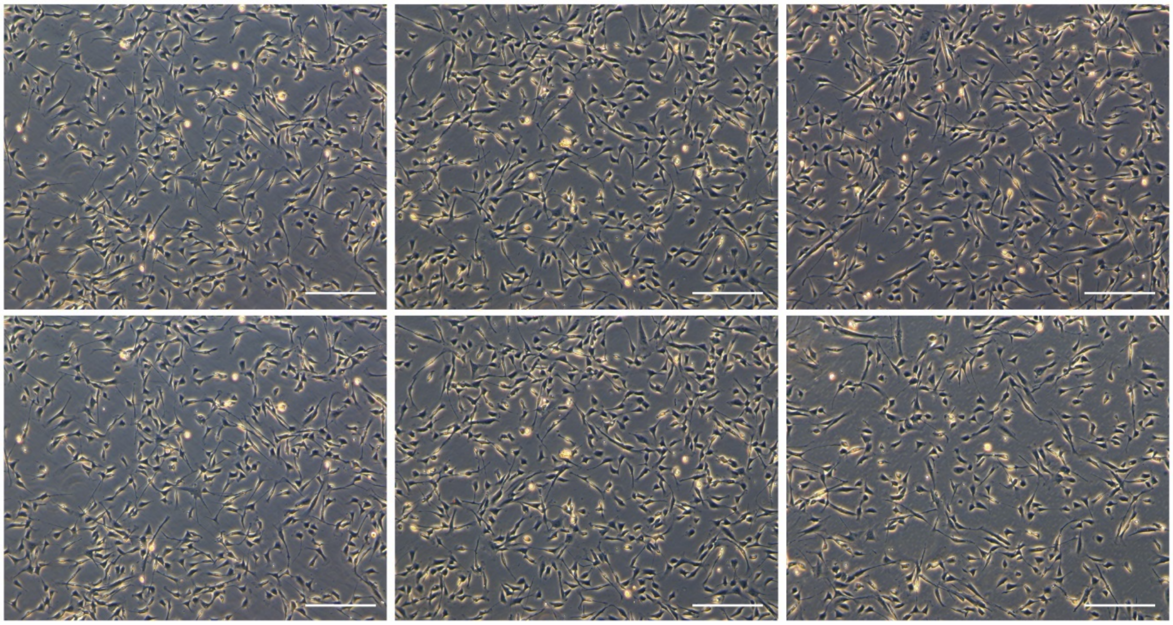


B

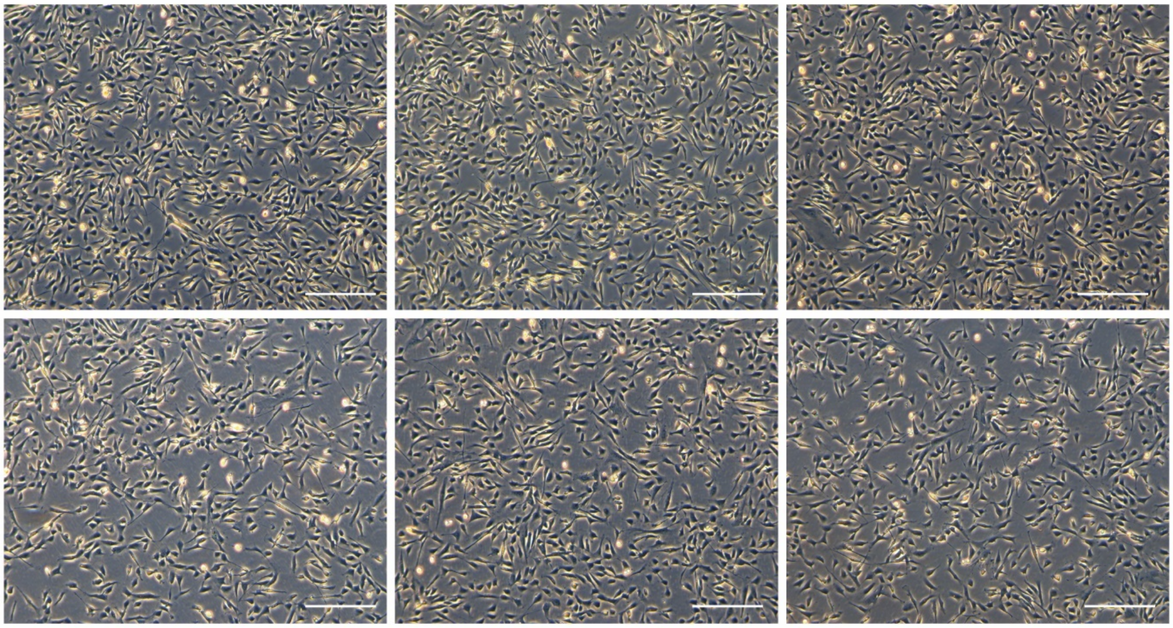


C

| GM | | GM1 | GM2 | GM3 |
|---|---|---|---|---|
| Doubling time | iBatM-Pmeso-S | 33.35 | 27.08 | 20.29 |
| | iBatM-Pmeso-TC | 24.37 | 20.29 | 23.33 |

**Figure S5. Comparison of Growth Media Effects on immortalized cells.** A. Phase-contrast images showing the morphology of self-immortalized myoblasts iBatM-Pmeso-S at passage 41 (P41) and P46 cultured in F10, DMEM/F10, or DMEM-based myoblast growth media. B. Phase-contrast images of hTERT/CDK4-immortalized myoblasts iBatM-Pmeso-TC at different passages cultured under the same media conditions. C. Quantification of doubling time for both iBatM-Pmeso-S and iBatM-Pmeso-TC in F10, 1:1 F10/DMEM, and DMEM, highlighting media-dependent differences in proliferative capacity. Scale bar: 100 μm.

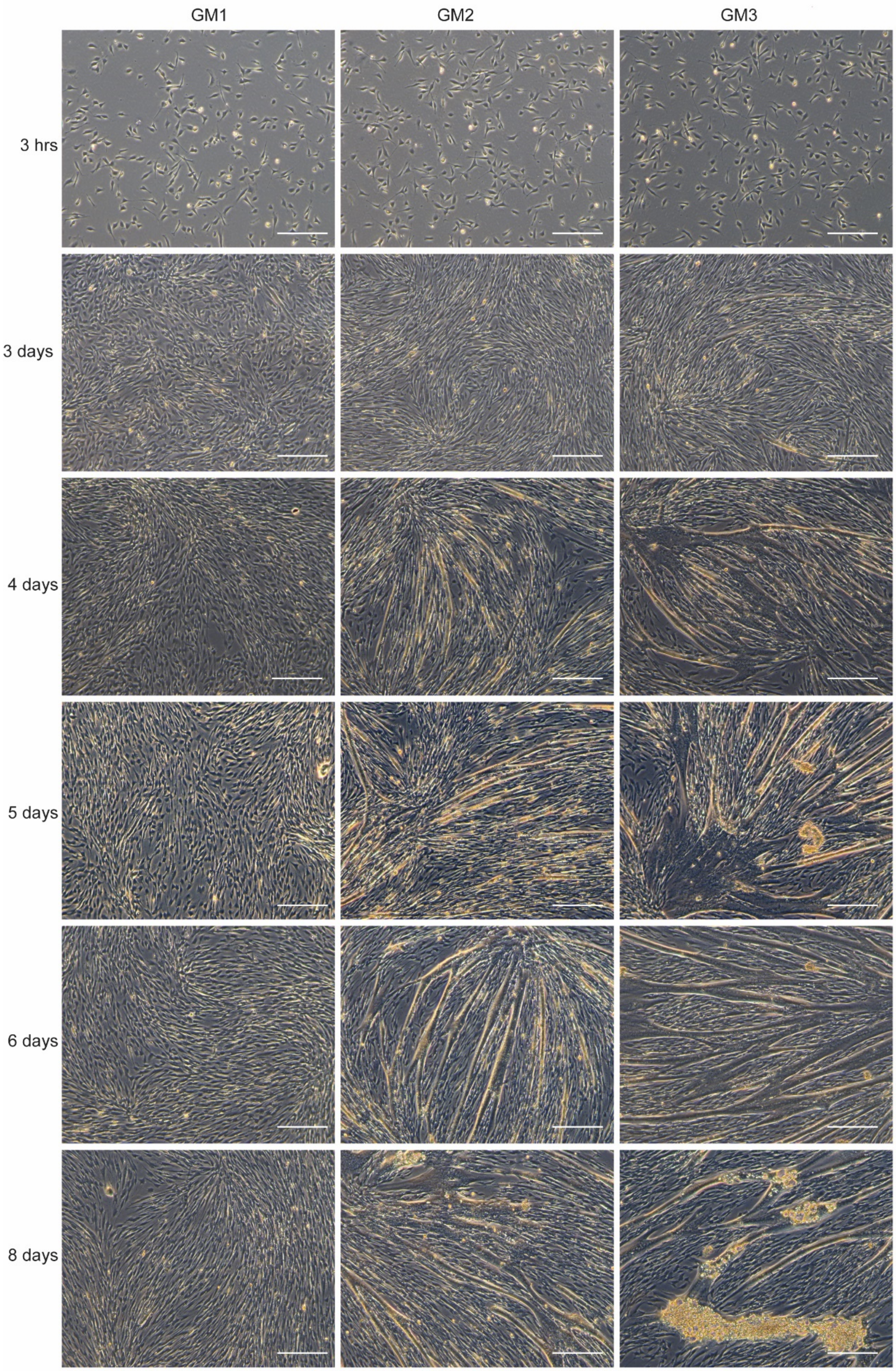

GM1
GM2
GM3
3 hrs
3 days
4 days
5 days
6 days
8 days

**Figure S6. DMEM Media Supports Myotube Formation in iBatM-Pmeso-S.** Phase-contrast images showing the morphological progression of iBatM-Pmeso-S cultured in GM1, GM2 and GM3 over an 8-day time course. While all media supported early attachment (3 hours) and proliferation (<3 days), only media containing DMEM promoted myotube differentiation by days 6-8. Scale bar: 100 μm.

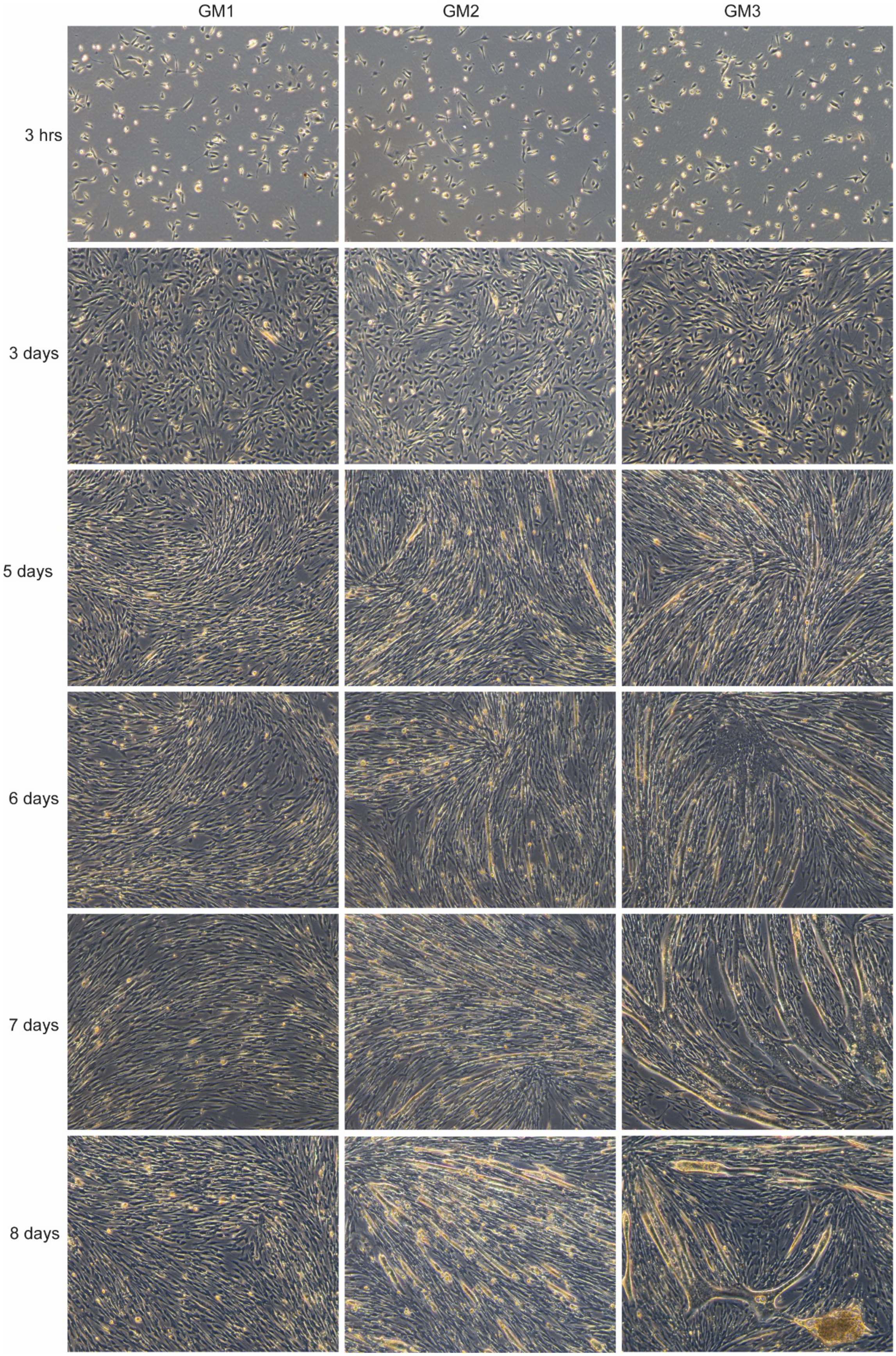
GM1
GM2
GM3
3 hrs
3 days
5 days
6 days
7 days
8 days

**Figure S7. DMEM Media Supports Myotube Formation in iBatM-Pmeso-TC.** Phase-contrast images showing the morphological progression of iBatM-Pmeso-TC cultured in GM1, GM2 and GM3 over an 8-day time course. While all media supported early attachment (3 hours) and proliferation (<3 days), only media containing DMEM promoted myotube differentiation by days 6-8. Scale bar: 100 μm.

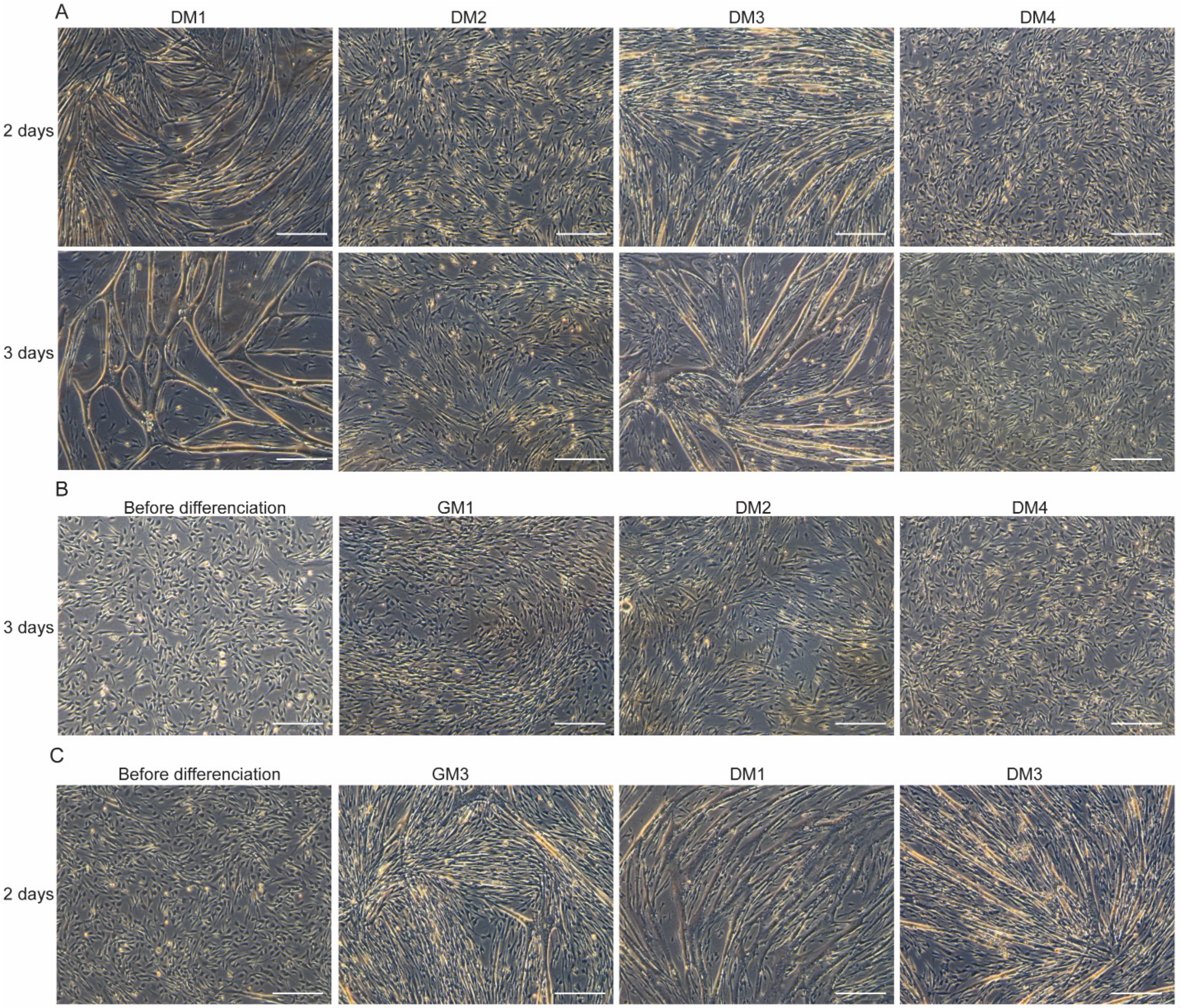


**Figure S8. Differentiation of iBatM-Pmeso-S Depend on Media Composition.** A. Phase-contrast images of self-immortalized myoblasts (iBatM-Pmeso-S) cultured for 2 and 3 days in four different media formulations (DM1-DM4; Table 2). Morphological differences reflect varying capacities of each media to support myotube formation. B. iBatM-Pmeso-S cultured after 3 days in GM1 versus DM2 and DM4. These conditions failed to support differentiation, as indicated by continued mononuclear morphology and lack of alignment or fusion. C. Comparisons of iBatM-Pmeso-S cultured in GM3, DM1, and DM3 after 2 days. Cell fusion, elongation, and alignment are observed. Scale bar: 100 µm.

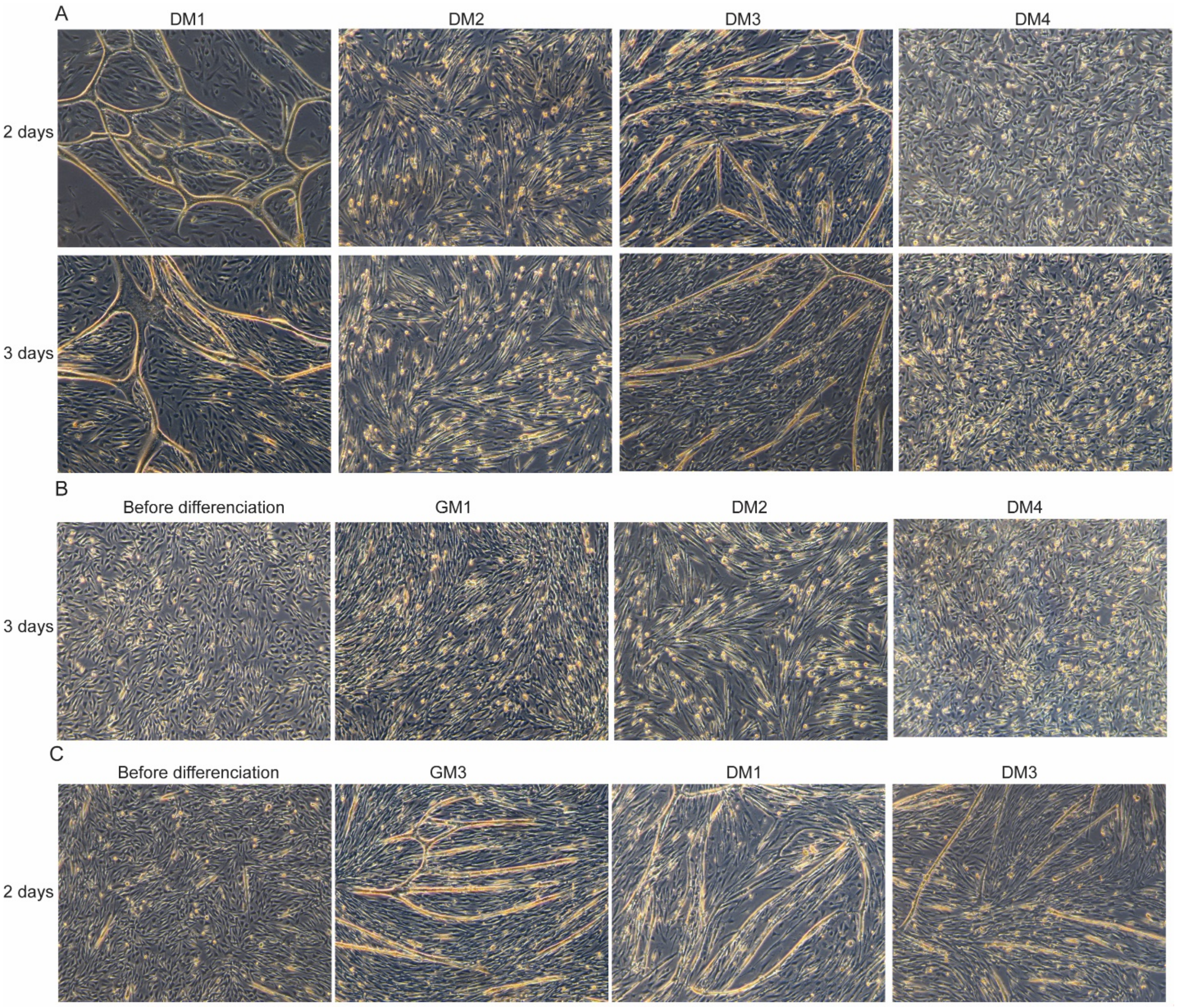


**Figure S9. Differentiation of iBatM-Pmeso-TC Depend on Media Composition.** A. Phase-contrast images of hTERT/CDK4-immortalized myoblasts (iBatM-Pmeso-TC) cultured for 2 and 3 days in four different media formulations (DM1-DM4; Table 2). B. Cells cultured in GM1, DM2, and DM4 for 3 days. These conditions fail to support clear myotube formation. **C.** 2 days after culturing in DMEM, DM1, and DM3, Cell fusion, elongation, and alignment are observed. Scale bar: 100 µm.
